# One Tracer, Dual Platforms: Unlocking Versatility of Fluorescent Probes in TR-FRET and NanoBRET Target Engagement Assays

**DOI:** 10.1101/2025.03.24.645143

**Authors:** Erika Y. Monroy, Xin Yu, Dong Lu, Xiaoli Qi, Jin Wang

## Abstract

Target engagement assays are essential for drug discovery, utilizing Time-Resolved Fluorescence Resonance Energy Transfer (TR-FRET) and Nano Bioluminescence Resonance Energy Transfer (NanoBRET) as complementary methods for biochemical and cellular evaluation. Traditional platforms require distinct fluorescent tracers, increasing costs and complexity. This study systematically evaluates the cross-platform performance of T2-BODIPY-FL and T2-BODIPY-589, tracers developed for receptor-interacting protein kinase 1 (RIPK1) target engagement in TR-FRET and NanoBRET applications, respectively. Our results demonstrate both tracers effectively bridge biochemical and cellular assays, providing reliable measurements. T2-BODIPY-589 demonstrates superior performance in NanoBRET (Z’ up to 0.80) and acceptable functionality in TR-FRET (Z’=0.53). Conversely, T2-BODIPY-FL performs optimally in TR-FRET (Z’=0.57) and exhibits NanoBRET potential (Z’ up to 0.72). Competition assays with an unlabeled inhibitor yielded consistent binding constants across all combinations. These findings suggest a single tracer can integrate diverse assay platforms, enhancing consistency and comparability in drug discovery.

**Table of Content Graphic:** 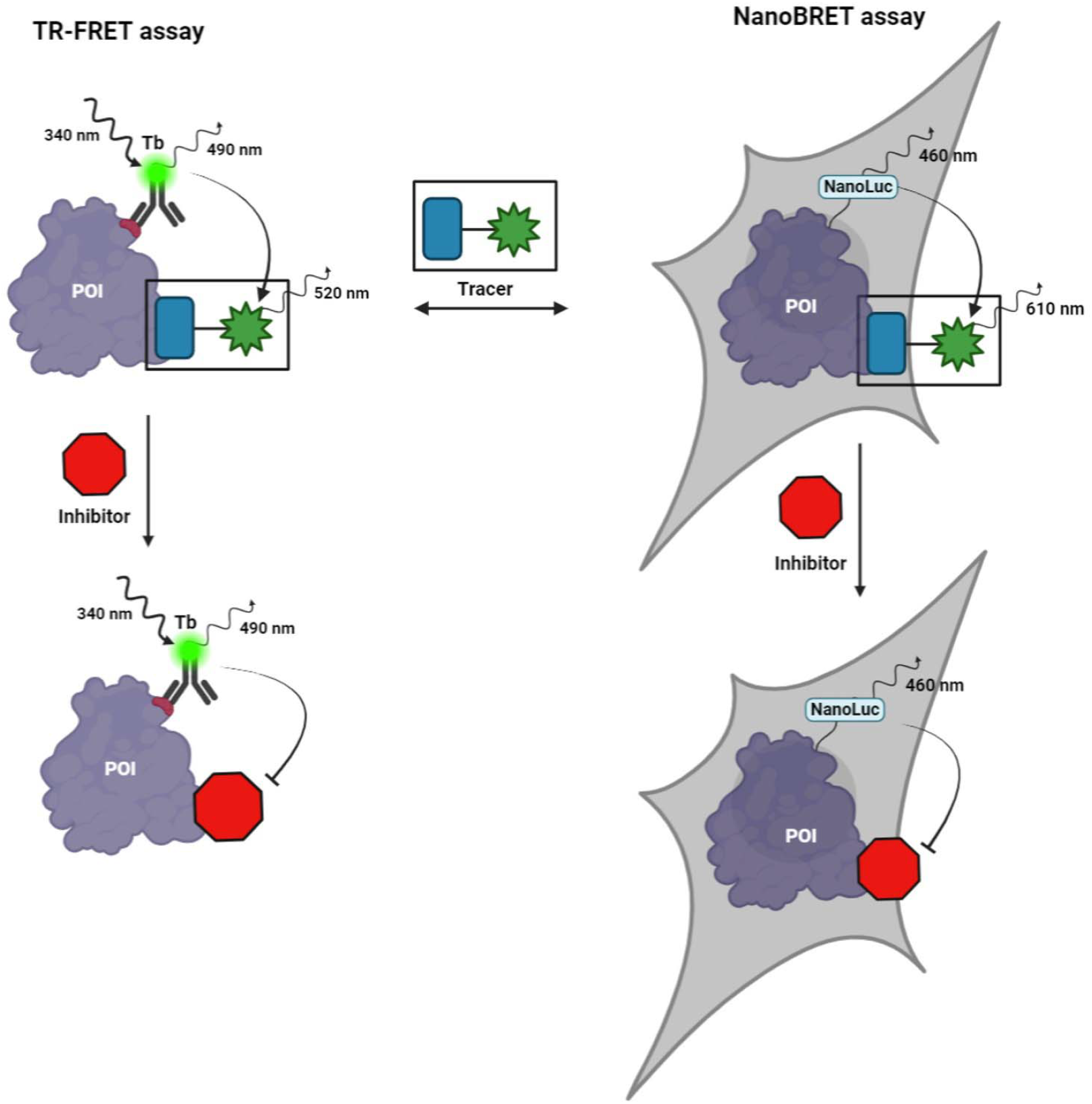

## INTRODUCTION

Biomedical research has accelerated significantly in recent decades, driven by advancements in biochemical, cellular, and tissue-based assays. These advancements expedite new therapeutic discovery by elucidating crucial protein-protein interactions for effective treatment strategies.^1^ While methods like surface plasmon resonance, thermal shift assays, nuclear magnetic resonance, and various calorimetry techniques measure direct target-compound interactions, many face limitations such as low throughput, protein purification requirements, or high noise level.^2–4^

Fluorescence/luminescence-based technologies are critical tools in modern molecular biology and drug discovery, particularly for high-throughput screening. Their advantages include high sensitivity, signal stability, adaptability across different assay formats, operational simplicity, and non-destructive analysis.^5–8^ Their versatility makes them indispensable across scientific disciplines, from detecting viral proteins in infected cells to screening diverse compound libraries against challenging therapeutic targets.^9,10^

Among these technologies, Time-Resolved Fluorescent Resonance Energy Transfer (TR-FRET) and Nano Bioluminescent Resonance Energy Transfer (NanoBRET) have gained prominence. Both noninvasive methods are mainstays in drug discovery for measuring protein-ligand interactions due to their high sensitivity and low background noise, leading to their growing popularity in preclinical biomedical research. ^11,12^ Assay performance assessment using hit confirmation criteria has been extensively studied by Zhang and colleagues and is summarized in Table S1.^13^

TR-FRET combines fluorescence sensitivity with FRET specificity to study molecular interactions. This technique relies on energy transfer between a donor (typically a lanthanide chelate) and an acceptor fluorophore when they are in close proximity (1-10 nm).^14^ TR-FRET utilizes donor fluorophores with exceptionally long fluorescent lifetimes (microseconds), enabling time-resolved detection that minimize background fluorescence and enhances signal-to-noise ratios.^14–16^ TR-FRET assays allow quantitative measurements of protein-ligand interactions, integrating seamlessly into high-throughput screening platforms for drug discovery.^17–20^

NanoBRET, developed by Promega Corporation, leverages bioluminescent resonance energy transfer to monitor target-ligand interactions in real time under physiologically relevant conditions.^21,22^ At its core, NanoBRET employs NanoLuc, a compact 19 kDa luciferase enzyme. NanoLuc emits light upon catalyzing its substrate, furimazine, generating a bioluminescent signal that excites nearby acceptor fluorophores within a critical 1-10 nm proximity range.^23^ Unlike TR-FRET, NanoBRET’s spontaneous light generation from enzymatic catalysis eliminates several technical challenges, minimizing autofluorescence and photobleaching while significantly reducing background noise. This technique is invaluable for tracking molecular interactions as they occur within the complex environment of living cells, providing insights into target engagement dynamics by small molecules and biologics under conditions that closely mirror the native cellular context.^24–29^

Fluorescent tracers are crucial components in both TR-FRET and NanoBRET assays, though their spectral requirements differ significantly between platforms.^30^ Terbium-based TR-FRET assays typically select fluorophores absorbing at 488 nm, leveraging the lanthanide donor’s characteristic emission bands.^31–33^ Conversely, NanoBRET assays use red-shifted fluorophores such as BODIPY 576/589 or NCT derivatives as acceptors. These are chosen for their minimal spectral overlap with NanoLuc donor emission to optimize energy transfer efficiency and reduce background signal.

Drug discovery campaigns often require both TR-FRET and NanoBRET platforms for comprehensive characterization of compound-target interactions from biochemical to cellular contexts. This dual requirement traditionally necessitates synthesizing separate tracers conjugated with platform-specific fluorophores, increasing development costs and potential variability between assay formats.

In this study, we evaluate the cross-platform compatibility of tracers originally designed for a single assay type. Specifically, we assess whether a TR-FRET-optimized tracer can effectively function in NanoBRET applications and vice versa. Using two fluorescent tracers, T2-BODIPY-FL (T2-BDP-FL) and T2-BODIPY-576/589 (T2-BDP-589), initially developed for RIPK1 target engagement studies, we systematically characterize their performance across both platforms under various detection parameters and assay conditions.^8,20,34,35^ These tracers demonstrate versatility across biochemical and cellular assay formats. Through comprehensive comparative analysis, we show that thoughtfully designed fluorescent tracers can bridge the gap between in vitro and cellular assay systems, delivering consistent and biologically relevant data while streamlining the tracer development process for multi-platform drug discovery applications.

For clarity, the BODIPY 576/589 (BDP589) dye used for tracer development in this work has the same chemical structure as Promega’s NanoBRET 590 (NB590).^36^ The difference is only in nomenclature.

## RESULTS

### Photophysical characterization

For comprehensive target engagement studies across biochemical and cellular platforms, we previously developed two specialized fluorescent tracers: T2-BDP-FL (Figure 1A) for TR-FRET and T2-BDP-589 (Figure 1B) for NanoBRET assays.^8^

**Figure 1.**
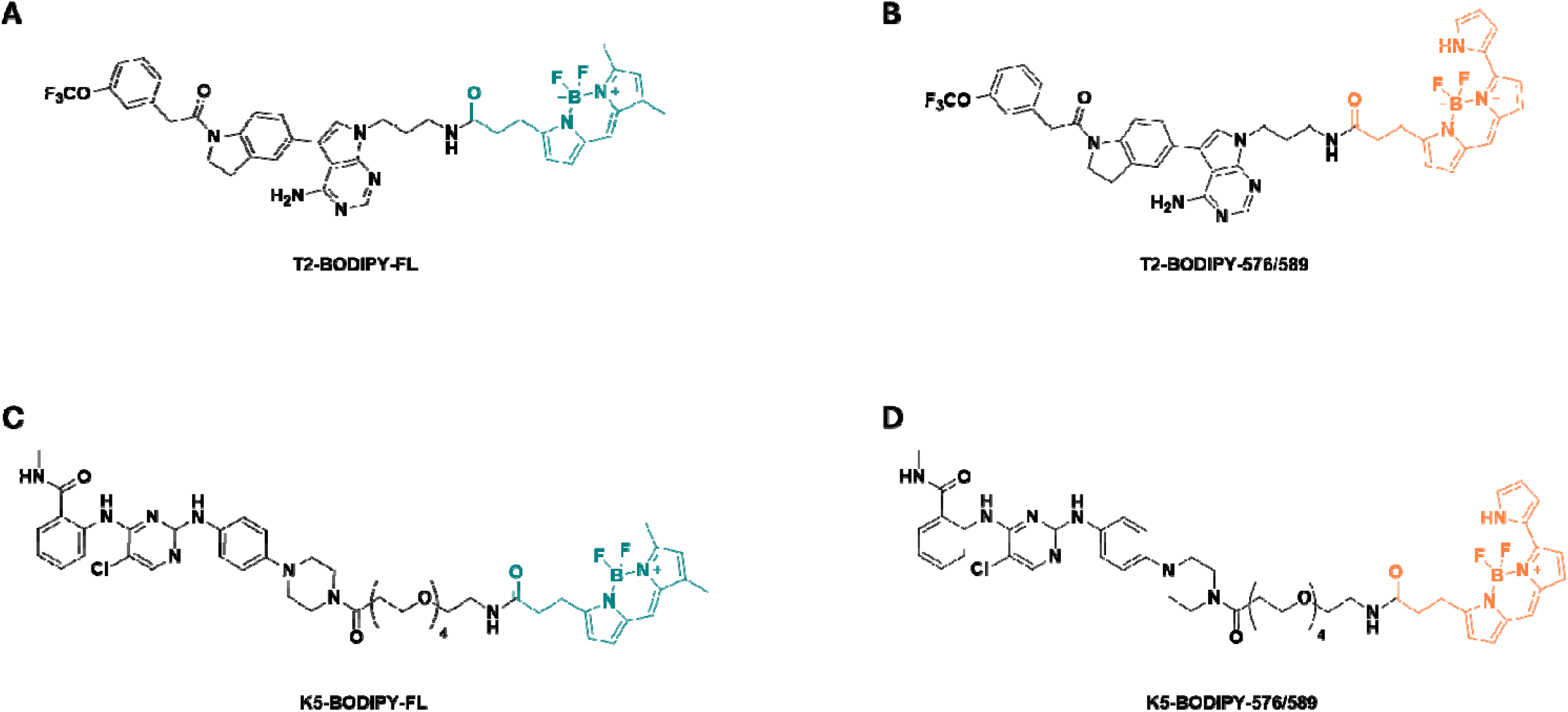
Chemical structures of fluorescent tracers used for cross-platform evaluation. (A) T2-BODIPY-FL (T2-BDP-FL) and (B) T2-BODIPY-576/589 (T2-BDP-589) were employed for RIPK1 target engagement assays. (C) K5-BODIPY-FL (K5-BDP-FL) and (D) K5-BODIPY-576/589 (K5-BDP-589) were used for BTK assays.

In TR-FRET assays, terbium (Tb) serves as an ideal donor due to its long fluorescence lifetime (milliseconds vs. nanoseconds) enabling time-gated detection and improved signal-to-noise ratios.^32,37^ Upon excitation at 340 nm, Tb exhibits four major emission transitions from ^5^D_4_ excited state to various ^7^F_J_ ground states (J = 6, 5, 4, 3), producing characteristic narrow emission bands at approximately 490, 548, 587, and 620 nm. The first transition (^5^D_4_→^7^F_6_) at 490 nm overlaps well with T2-BDP-FL’s absorption spectrum (Figure 2A), ensuring high energy transfer efficiency.^38^ Conventional TR-FRET assays with this pair typically use 520 ± 10 nm/490 ± 10 nm filter settings, capturing T2-BDP-FL emission while effectively discriminating against the Tb signal.

**Figure 2.**
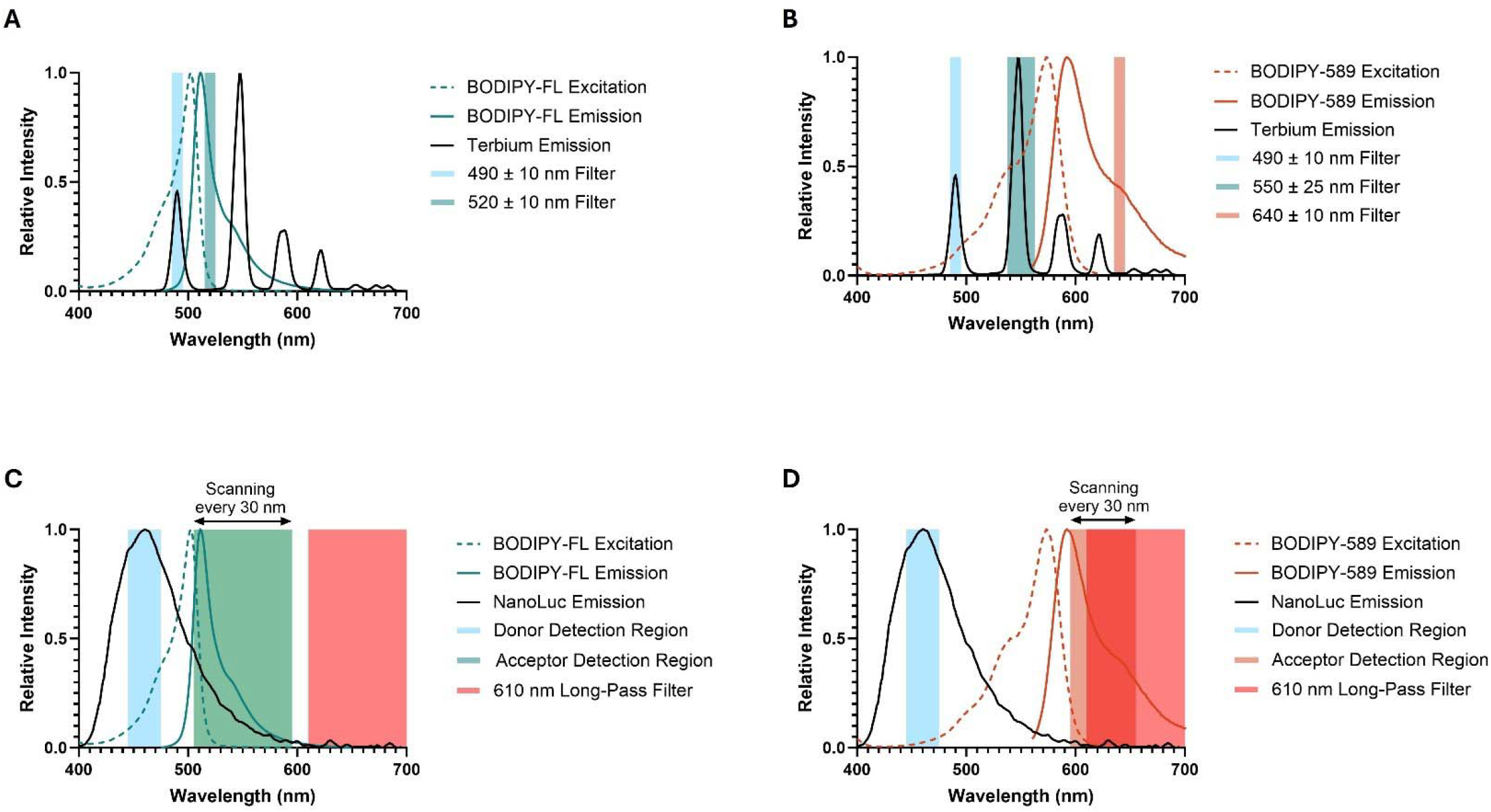
Spectral characterization of fluorescent tracers with TR-FRET and NanoBRET donors and optimized detection windows. (A) BODIPY-FL excitation and emission spectra with Terbium (Tb) emission and corresponding filter settings for TR-FRET. (B) BODIPY-589 excitation and emission spectra with Terbium emission and corresponding filter settings for TR-FRET. (C) BODIPY-FL excitation and emission spectra with NanoLuc emission and detection windows for NanoBRET. (D) BODIPY-589 excitation and emission spectra with NanoLuc emission and detection windows for NanoBRET.

Interestingly, Tb’s second (^5^D_4_→^7^F_5_ at 548 nm) and third (^5^D_4_→^7^F_4_ at 587 nm) transitions also overlap with T2-BDP-589’s absorption spectrum (Figure 2B).^39^ This compatibility suggests T2-BDP-589’s potential versatility in TR-FRET assays, optimizable with alternative filter settings like 640 ± 10 nm/490 ± 10 nm or 640 ± 10 nm/550 ± 25 nm to capture its red-shifted emission while maintaining Tb discrimination.

In NanoBRET assays, NanoLuc luciferase generates a blue luminescent signal with a maximum emission at 460 nm, tapering to approximately 600 nm.^40^ T2-BDP-FL’s absorption spectrum significant overlap with this NanoLuc emission profile (Figure 2C), theoretically enabling energy transfer in cellular systems. However, this spectral proximity can cause signal bleed-through from the intense donor into the acceptor channel. To address this, we evaluated various emission settings using a linear variable filter (LVF) monochromator detector, incrementally adjusted in 30 nm bandwidths, alongside a 610 nm long-pass filter. This identified optimal detection parameters maximizing T2-BDP-FL signal while minimizing NanoLuc bleed-through. Similarly, we assessed T2-BDP-589’s performance (Figure 2D), which was designed with red-shifted spectral properties to enhance separation from the NanoLuc emission and reduce cellular assay interference.

This comprehensive photophysical characterization of both tracers across multiple detection parameters aimed to determine their versatility across TR-FRET and NanoBRET platforms, potentially simplifying assay development and increasing experimental flexibility in target engagement studies.

### Optimizing TR-FRET: Cross-Platform Utility of T2-BDP-FL and T2-BDP-589 for RIPK1 Target Engagement

To evaluate the cross-platform compatibility, we optimized TR-FRET assay conditions using T2-BDP-FL and T2-BDP-589 with His-tagged RIPK1 and an anti-6xHis-Tb antibody. Three filter pair combinations (520 ± 10 nm/490 ± 10 nm, 640 ± 10 nm/490 ± 10 nm, and 640 ± 10 nm/550 ± 25 nm) were tested to determine optimal detection parameters.

Results, summarized in Table S2, revealed distinct performance profiles. T2-BDP-FL performed best with the 520 ± 10 nm/490 ± 10 nm filter pair (Z’ = 0.57, signal window = 6.0), consistent with its spectral properties and TR-FRET compatibility, but showed negative Z’ values with other combinations. T2-BDP-589, designed primarily for NanoBRET, demonstrated adequate TR-FRET performance with the 640 ± 10 nm/490 ± 10 nm filter pair (Z’ = 0.53, signal window = 7.8) and acceptable performance with the 520 ± 10 nm/490 ± 10 nm filter pair (Z’ = 0.40, signal window = 10.5). However, its performance was marginal with the 640 ± 10 nm/550 ± 25 nm filter pair (Z’ = 0.29, signal window = 3.1).

Dose-response analysis (Figures 3A to 3C) confirmed both tracers’ functionality in TR-FRET. T2-BDP-FL exhibited an equilibrium dissociation constant (K_d_) of 608 nM with the 520 ± 10 nm/490 ± 10 nm filter pair, while T2-BDP-589 showed a K_d_ of 443 nM with the 640 ± 10 nm/490 ± 10 nm filter pair. Although other filters (520 ± 10 nm/490 ± 10 nm and 640 ± 10 nm/550 ± 25 nm) met minimal assay feasibility, only conditions with Z’ values exceeding 0.5 were suitable for subsequent IC_50_ determination with unlabeled T2 inhibitor. This optimization establishes a robust and sensitive TR-FRET assay for RIPK1, ensuring reliable characterization of binders.

**Figure 3.**
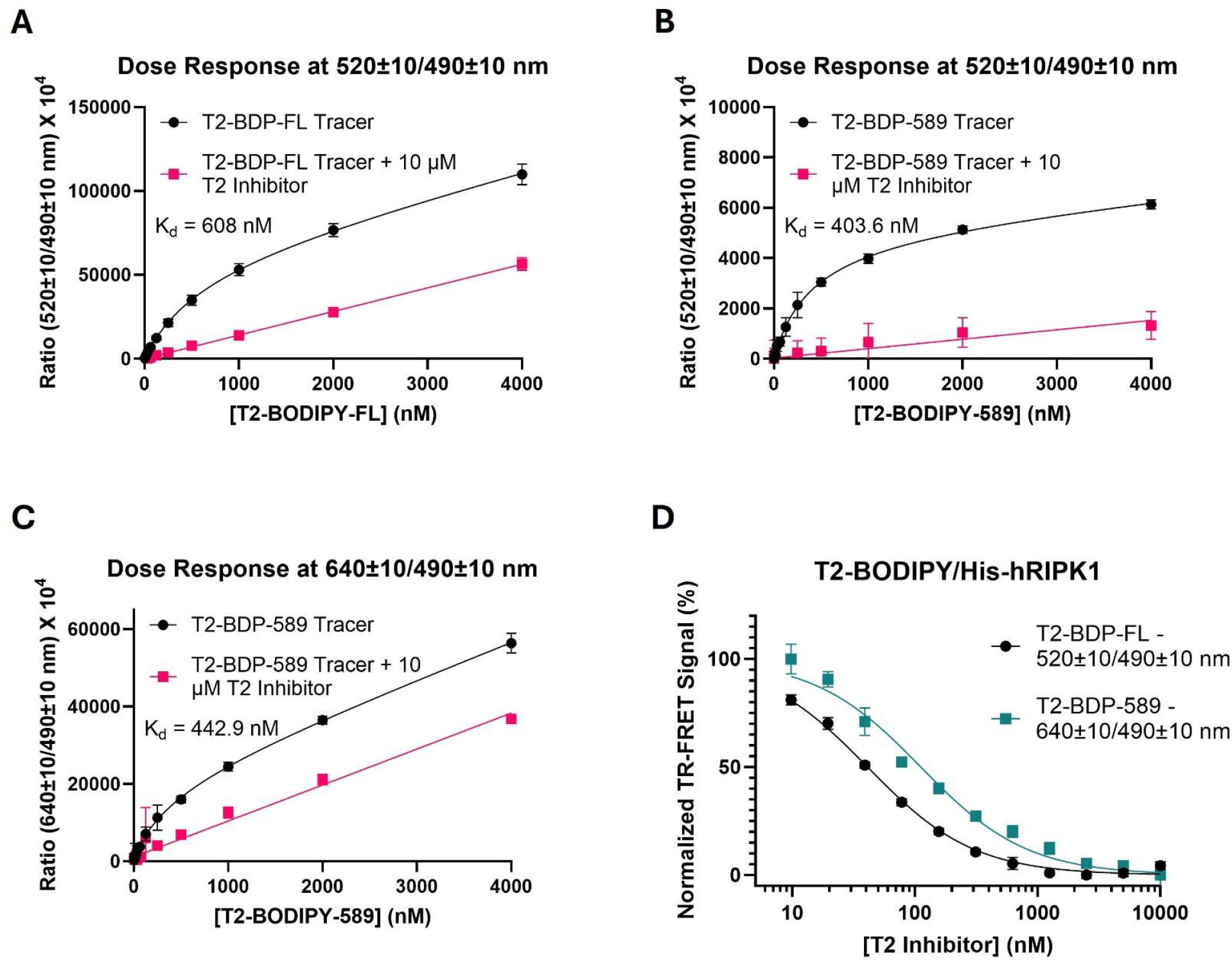
Optimization of TR-FRET assay for RIPK1 target engagement. (A-C) Saturation binding and nonspecific binding curves for 1 nM His-hRIPK1 enzyme with increasing concentrations of T2-BODIPY-FL (A), T2-BODIPY-589 (B, C) under different TR-FRET filter combinations. (D) Competition binding curves of unlabeled T2 inhibitor against His-hRIPK1 using optimized concentrations of T2-BODIPY-FL and T2-BODIPY-589 in TR-FRET assays. Data represent mean ± SD (n=6 technical replicates).

In Figure 3D, we apply optimized filter settings to assess the unlabeled T2 inhibitor’s relative affinity, a highly selective RIPK1 inhibitor. To accurately determine the inhibitory constant (K_i_), we selected fluorescent tracer concentrations at their K_d_ values using the Cheng-Prusoff equation (K_i_ = IC_50_/2).^41^ Interestingly, competition assays with the unlabeled T2 inhibitor (Table S3) yielded K_i_ values of 18 ± 0.70 nM for T2-BDP-FL and 54 ± 5.9 nM for T2-BDP-589. This discrepancy likely arises from differences in detection parameters rather than true binding affinity variations, as filter settings capture distinct emission regions, affecting signal intensity and binding metrics. Both tracers were effectively displaced by the unlabeled T2 inhibitor, demonstrating specific binding to RIPK1.

While the T2-BDP-589 probe allows for two possible filter settings, the T2-BDP-FL filter settings (520 ± 10 nm/490 ± 10 nm) better aligned with the TR-FRET assay’s emission spectrum, providing a more sensitive measure of unlabeled T2 inhibitory activity and ensuring greater consistency. These findings confirm T2-BDP-FL as the optimal choice for TR-FRET applications, while T2-BDP-589 demonstrates versatility and potential for cross-platform use with appropriate detection parameters. The generalizability of these optimized TR-FRET conditions was further supported by studies with Bruton’s tyrosine kinase (BTK), detailed in Supplemental Figures S1-S2 and Tables S4-S5.

### Cross-Platform Tracer Versatility: T2-BDP-FL and T2-BDP-589 in NanoBRET RIPK1 Target Engagement

To assess target versatility in cellular contexts, we evaluated T2-BDP-FL and T2-BDP-589 in NanoBRET assays using NanoLuc-tagged RIPK1 expressed in HEK293T cells. We employed an LVF monochromator detector to investigate various emission combinations and establish optimal detection parameters, adjusting the monochromator 30 nm bandwidth intervals (Table S6).

T2-BDP-589 demonstrated strong performance across various emission pairs using LVF monochromator and filters optical configurations. The highest Z’ factor (0.80) and a signal window (15) were achieved with the 610 nm long-pass filter and the 460 ± 30 nm monochromator setting. The 610 ± 30 nm and 460 ± 30 nm settings yielded a Z’ factor of 0.79 and a signal window of 17. In contrast, the 640 ± 30 nm and 460 ± 30 nm combination resulted in a Z’ factor of 0.58 and a signal window of 7.7, while the 580 ± 30 nm and 460 ± 30 nm setting yielded a Z’ factor of 0.62 and a signal window of 7.6. These results confirm T2-BDP-589 is well-suited for NanoBRET applications, exhibiting robust assay performance across multiple detection parameters.

Despite being optimized for TR-FRET, T2-BDP-FL also demonstrated effective NanoBRET compatibility, particularly with the 520 ± 30 nm and 460 ± 30 nm monochromator settings (Z’ = 0.72, signal window = 14). However, performance declined with the 610 nm long-pass filter and the 460 ± 30 nm setting (Z’ = 0.21, signal window = 3.1), likely due to insufficient emission from T2-BDP-FL at longer wavelengths.

Dose-response analysis provided further insights into tracer binding affinities. T2-BDP-589 exhibited K_d_ values ranging from 232 nM to 267 nM across different monochromator and filter combinations (Figures 4D-F), demonstrating strong and specific binding. T2-BDP-FL displayed slightly higher K_d_ values, ranging from 619 nM to 711 nM (Figures 4A-C). While both tracers showed reduced performance when shifting the acceptor wavelength, T2-BDP-589 maintained superior overall reliability and assay performance. These optimizations are essential for accurate target engagement quantification, facilitating reliable NanoBRET assay results.

**Figure 4.**
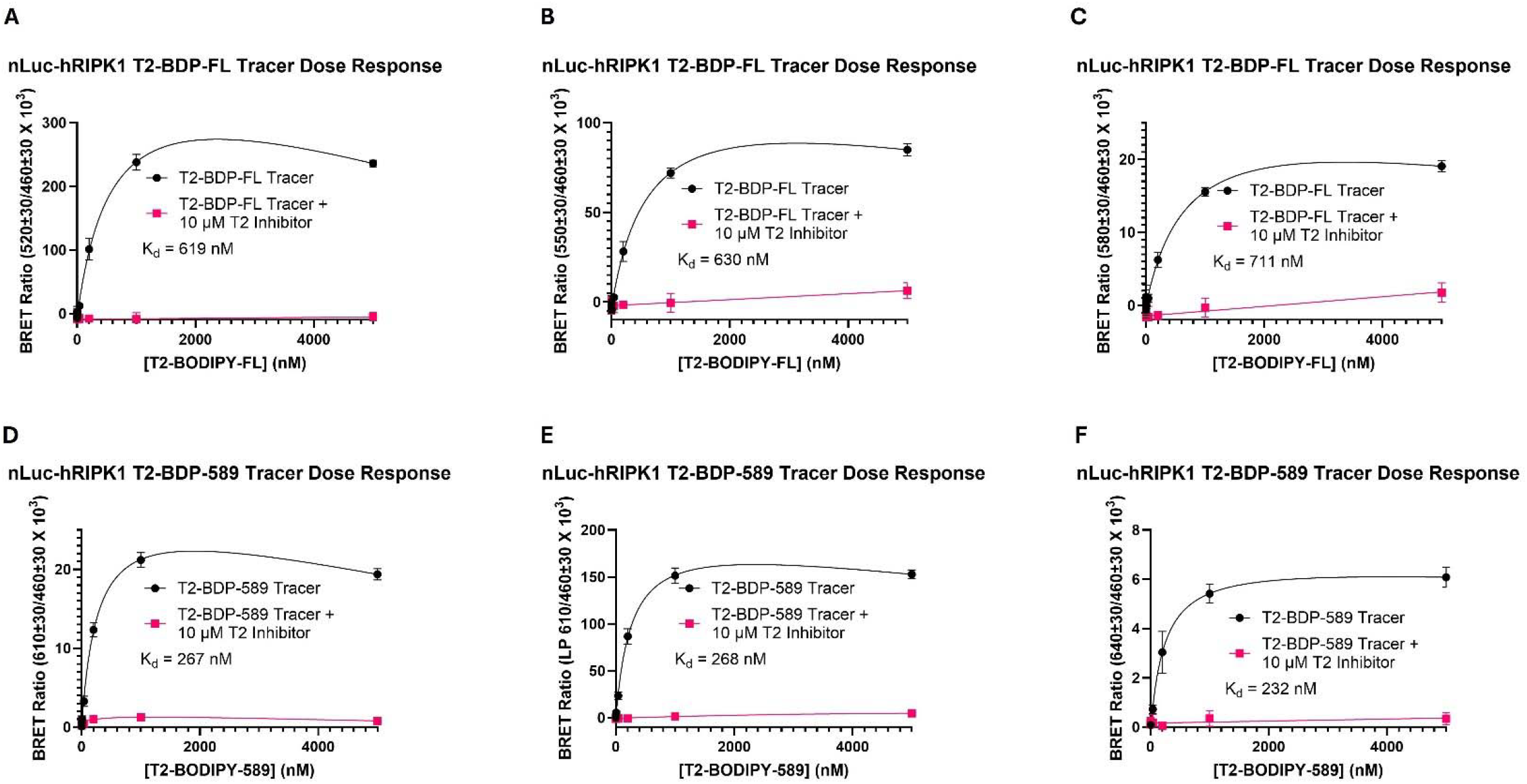
Validation of Tracer Compatibility for Enhanced NanoBRET Assay Performance. (A–C) Binding and nonspecific binding interactions of T2-BODIPY-FL to nLuc-hRIPK1 in HEK293T cells under different optical configuration settings: 520 ± 30 nm/460 ± 30 nm (A), 550 ± 30 nm/460 ± 30 nm (B), and 580 ± 30 nm/460 ± 30 nm (C). (D–F) Total and nonspecific binding of T2-BODIPY-589 to nLuc-hRIPK1 in HEK293T cells with optical configuration settings of 610 ± 30 nm/460 ± 30 nm (D), 610 nm LP/460 ± 30 nm (E), and 640 ± 30 nm/460 ± 30 nm (F), illustrating the effect of varying concentrations on binding efficacy. Data in A–F are presented as mean ± SD (n = 4 technical replicates).

Competition assays with the unlabeled T2 inhibitor (Figure 5 and Table S7) yielded consistent K_i_ values across all monochromator and filter combinations (6.6 ± 1.6 nM to 14 ± 2.5 nM). Minimal variation in K_i_ values with different tracers or detection settings indicated robust and reliable assay performance regardless of detection configuration. These findings demonstrate both tracers’ remarkable versatility in NanoBRET assays with appropriate wavelength combinations. T2-BDP-589 performs optimally in this format, requiring lower probe concentrations for effective target engagement, enhanced signal specificity, reduced risk of inner-filter effects, and minimal probe-induced cytotoxicity. The effectiveness of T2-BDP-FL in NanoBRET highlights the potential for using a single tracer across multiple assay platforms, streamlining assay development and enabling direct comparison of biochemical and cellular data. The versatility and scalability of this system were further reinforced by successful NanoBRET target engagement assays in HeLa cells with BTK-NanoLuc, as detailed in Supplemental Figures S3-S4 and Tables S8-S9.

**Figure 5.**
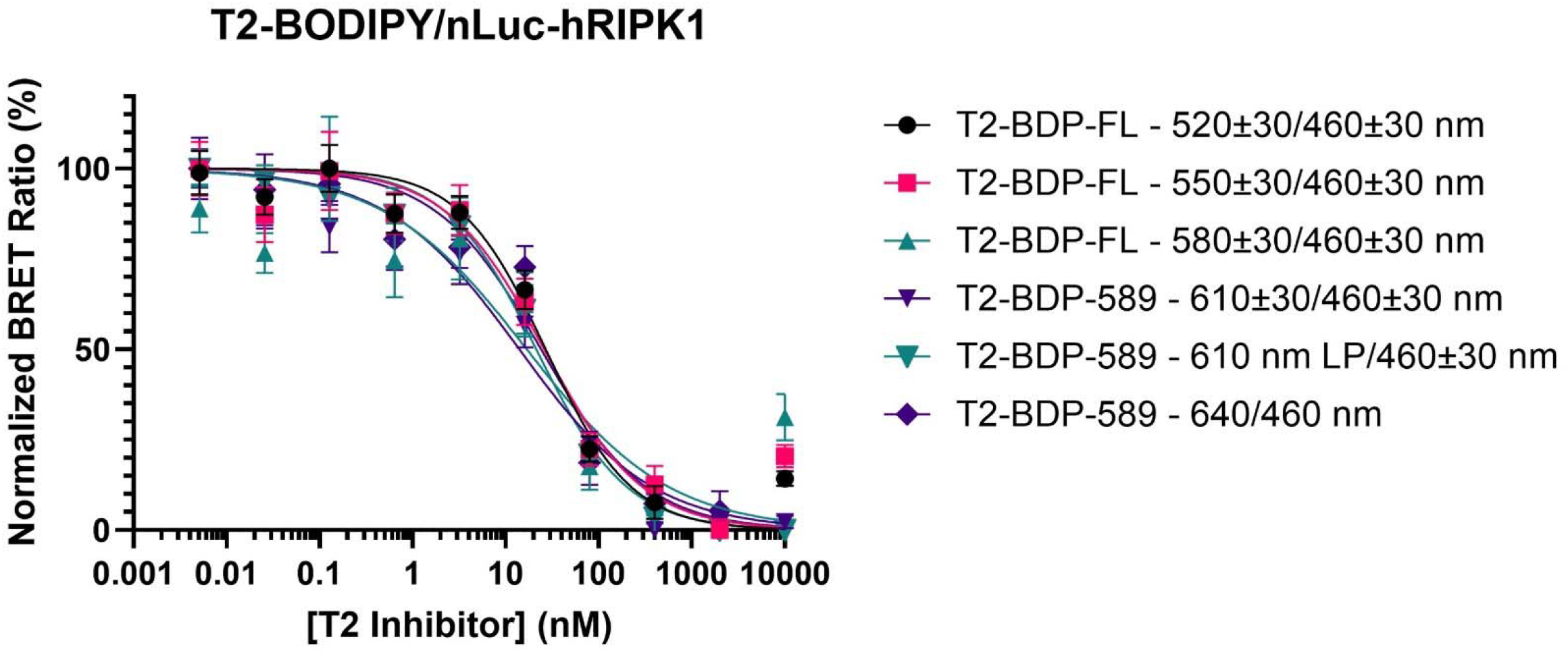
NanoBRET competition assays for RIPK1 target engagement. Normalized BRET ratio curves demonstrate displacement of T2-BODIPY-FL and T2-BODIPY-589 tracers by increasing concentrations of unlabeled T2 inhibitor in NanoLuc-RIPK1 expressing HEK293T cells, across various NanoBRET detection parameters. Data are presented as mean ± SD (n=4 technical replicates).

## DISCUSSION

Our evaluation of T2-BDP-FL and T2-BDP-589 across TR-FRET and NanoBRET platforms reveals unexpected tracer versatility, challenging the conventional approach of developing platform-specific probes.

Both assays determined the binding affinities of a T2 inhibitor against RIPK1, a therapeutic target. In TR-FRET assays, T2-BDP-FL exhibited a K_d_ of 608 nM (Z’ = 0.57, signal window = 6.0) with the 520 ± 10 nm/490 ± 10 nm filter pair. T2-BDP-589 demonstrated a slightly lower K_d_ of 443 nM (Z’ = 0.53, signal window = 7.8) with the 640 ± 10 nm/490 ± 10 nm filter pair. This suggests T2-BDP-589 may have a higher affinity under optimized TR-FRET conditions. However, K_d_ variations may stem from filter settings capturing distinct emission regions, affecting signal intensity and binding metrics, rather than true affinity differences. Competition assays with an unlabeled T2 inhibitor yielded different K_i_ values (18 ± 0.70 nM for T2-BDP-FL and 54 ± 5.9 nM for T2-BDP-589), with discrepancies likely arising from detection parameters differences.In NanoBRET assays, both tracers demonstrated strong binding to RIPK1, with T2-BDP-589 outperforming T2-BDP-FL, achieving the highest Z’ factor (0.80) and signal window (15) using the 610 nm long-pass filter and 460 ± 30 nm monochromator setting. This improved performance is attributed to greater spectral separation between NanoLuc and T2-BDP-589, reducing inner-filter effects and signal bleed-through, ensuring more reliable data. Surprisingly, T2-BDP-FL also showed effective NanoBRET compatibility, particularly with the 520 ± 30 nm and 460 ± 30 nm monochromator setting (Z’ = 0.72, signal window = 14), though its performance declined at longer wavelengths. Notably, similar K_i_ values (6.6-14 nM) across multiple configurations confirmed both tracers’ suitability for quantitative target engagement studies.

The cross-platform utility of these tracers was additionally validated across different kinase targets, including BTK, in both biochemical and cellular contexts (Figures S1-S2, Tables S4-S5 for TR-FRET; Figures S3-S4, Table S8-S9 for NanoBRET).T2-BDP-FLoffers a higher quantum yield and established performance in TR-FRET assays, making it excellent for biochemical screening.^42^ However, its shorter emission wavelength increases susceptibility to autofluorescent interference. In contrast, T2-BDP-589, red-shifted spectral properties provide superior resistance to such interference, especially valuable when screening compound libraries or working in complex biological matrices. Its longer emission wavelength (589 nm) enhances separation from both terbium and NanoLuc emissions, reducing signal bleed-through and improving signal-to-background ratios.

We recommend T2-BDP-589 as the preferred tracer for integrated assay development across both platforms. Its TR-FRET functionality, excellent performance in NanoBRET, and red-shifted spectral properties that minimize autofluorescent interference make it the more versatile choice for comprehensive target engagement studies. However, for laboratories focused exclusively on TR-FRET screening without cellular validation, T2-BDP-FL remains an excellent option.

The cross-platform compatibility of these fluorescent tracers represents a significant advancement with for assay development in drug discovery. This approach challenges the conventional notion that different assay platforms require distinct tracers, potentially streamlining probe development and reducing costs and complexity. By using identical tracers across biochemical and cellular formats, researchers can make more reliable comparisons between in vitro and cellular data without introducing tracer-specific variables that might otherwise confound interpretation. The principles established here have broad applicability beyond kinases, potentially extending to diverse target classes including membrane receptors, nuclear receptors, and protein-protein interactions, where consistent target engagement measurements across platforms provide valuable insights. Furthermore, this methodology lays the foundation for developing multiplexed assays capable of simultaneously monitoring interactions with multiple targets, offering a more comprehensive view of compound selectivity and engagement profiles.

In conclusion, our evaluation demonstrates that carefully designed fluorescent tracers can effectively function across both biochemical and cellular assay platforms. By selecting tracers with appropriate spectral properties, particularly those with red-shifted emission profiles like T2-BDP-589, researchers can streamline assay development, reduce variability, and obtain more consistent target engagement data across experimental systems.

## ASSOCIATED CONTENT

Supporting Information is available free of charge at the ACS Publications website.

- This includes additional references, comprehensive experimental methods detailing chemical synthesis and biological assays, as well as supplementary figures and tables (PDF)

## AUTHOR INFORMATION

### Author Contributions

E.Y.M. and J.W. designed experiments. E.Y.M performed experiments X.Y. and D.L. provided fluorescent tracers. E.Y.M. X.Q. and J.W. wrote the manuscript.

## ACKNOWLEDGEMENT

The research was supported in part by National Institute of Health (R01-CA268518 and R01-CA250503 to J.W.), Cancer Prevention & Research Institute of Texas (CPRIT, RP220480 to J.W.), and the Michael E. DeBakey, M.D., Professorship in Pharmacology to J.W. We also acknowledge Dr. Mazitschek and BMG Labtech’s advice and assistance with the customized filters.

## ABBREVIATIONS

TR-FRET: Time-Resolved Fluorescence Resonance Energy Transfer
NanoBRET: Nano Bioluminescence Resonance Energy Transfer
NanoLuc: NanoLuc luciferase
RIPK1: receptor-interacting protein kinase 1
HEK293T: human embryonic kidney 293T cells
K_d_: equilibrium dissociation constant
K_i_: inhibition constant
Z’: Z-prime factor
LP: longpass filter
BODIPY: dipyrrometheneboron difluoride
Tb: terbium
FL: fluorescein
NCT: nonchloro TOM
IC_50_: half-maximal inhibitory concentration
mM: millimolar
nm: nanometer
μM: micromolar
nM: nanomolar

