## Supplemental Info for "One Tracer, Dual Platforms: Unlocking Versatility of Fluorescent Probes in TR-FRET and NanoBRET Target Engagement Assays"

### Supporting Information

*Figure S1. TR-FRET saturation binding curves for BTK with fluorescent K5 tracers*

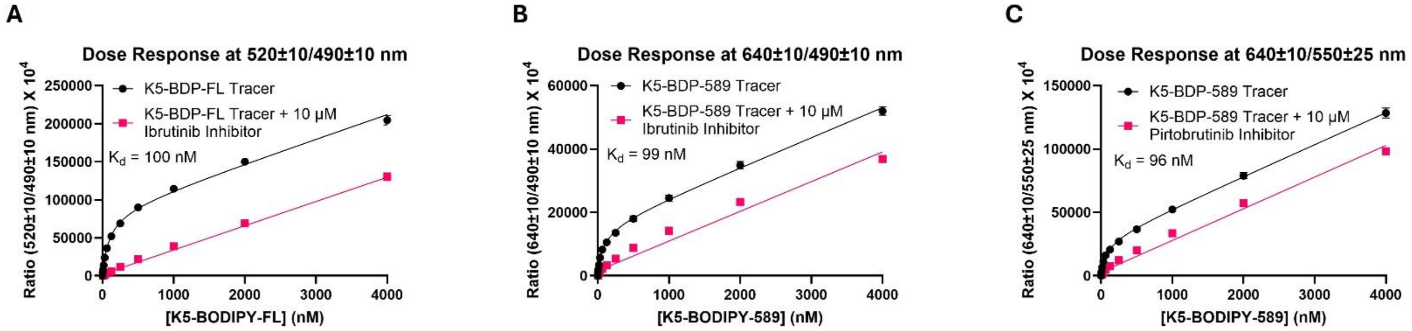

(A-C) Representative saturation binding curves for GST-BTK with increasing concentrations of K5-BODIPY-FL (A) and K5-BODIPY-589 (B, C) under different TR-FRET filter combinations. Data represent mean ± SD (n=6 technical replicates).

***Figure S2. TR-FRET competition assays for BTK target engagement with Ibrutinib*** [a]

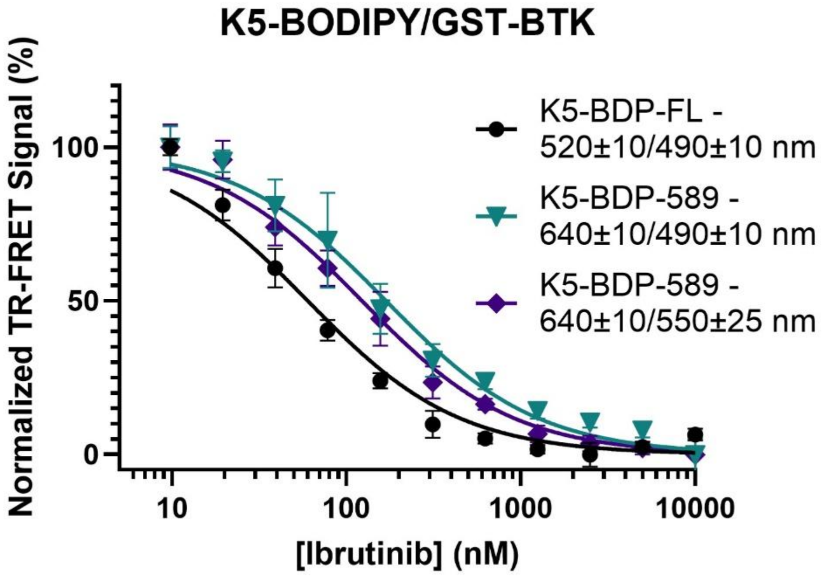

[a]Normalized TR-FRET signal curves demonstrating displacement of K5-BODIPY-FL and K5-BODIPY-589 tracers by increasing concentrations of Ibrutinib in GST-BTK biochemical assays, under various TR-FRET detection parameters. Data represent mean ± SD. (n=5 technical replicates).

***Figure S3. Representative saturation binding curves for K5 fluorescent tracers.***

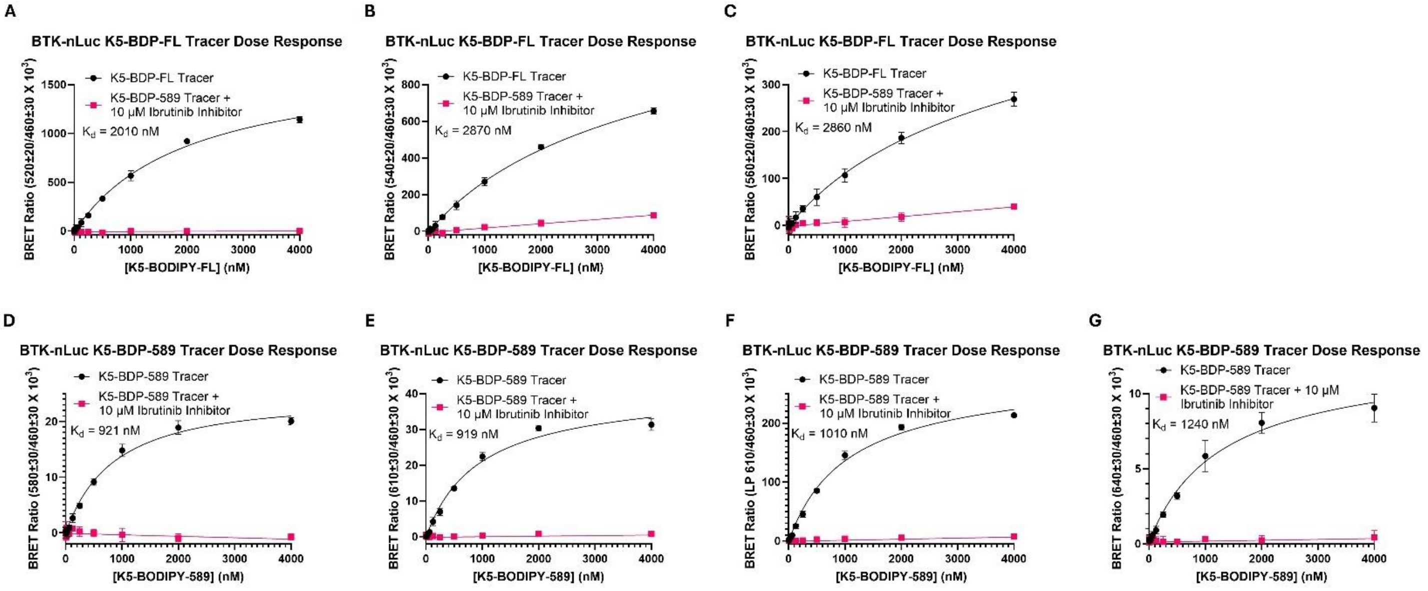

(A–C) Binding and nonspecific binding interactions of K5-BODIPY-FL to BTK-nLuc in HeLa cells under different optical configuration settings: 520 ± 20 nm/460 ± 30 nm (A), 540 ± 20 nm/460 ± 30 nm (B), and 560 ± 20 nm/460 ± 30 nm (C). (D–G) Total and nonspecific binding of K5-BODIPY-589 to BTK-nLuc in HeLa cells under different optical configuration settings of 580 ± 30 nm/460 ± 30 nm (D), 610 ± 30 nm/460 ± 30 nm (E), LP 610 nm/460 ± 30 nm (F), and 640 ± 30 nm/460 ± 30 nm (G), illustrating the effect of varying concentrations on binding efficacy. Data are presented as mean ± SD (n = 4 technical replicates).

***Figure S4. Assessment of Intracellular BTK Target Engagement of Ibrutinib Inhibitor Using NanoBRET Displacement Assay with Fluorescent Tracers [a]***

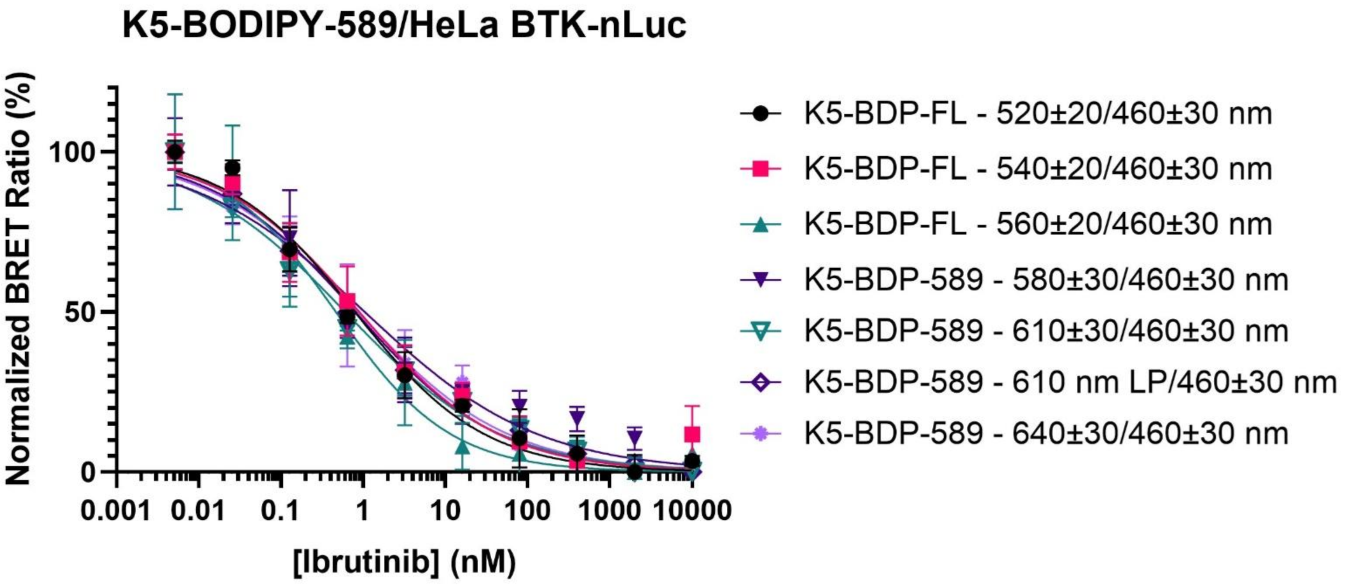

[a]Dose-response titration of ibrutinib inhibitor with transiently transfected BTK-nLuc in HeLa using K5-BODIPY-FL and K5-BDP-589 as fluorescent tracers. Data are presented as mean ± SEM (n=4 technical replicates).

*Figure S5. Time-Dependent Degradation of T2-BDP-FL Tracer in Cell Medium Monitored by LC-UV* [a]

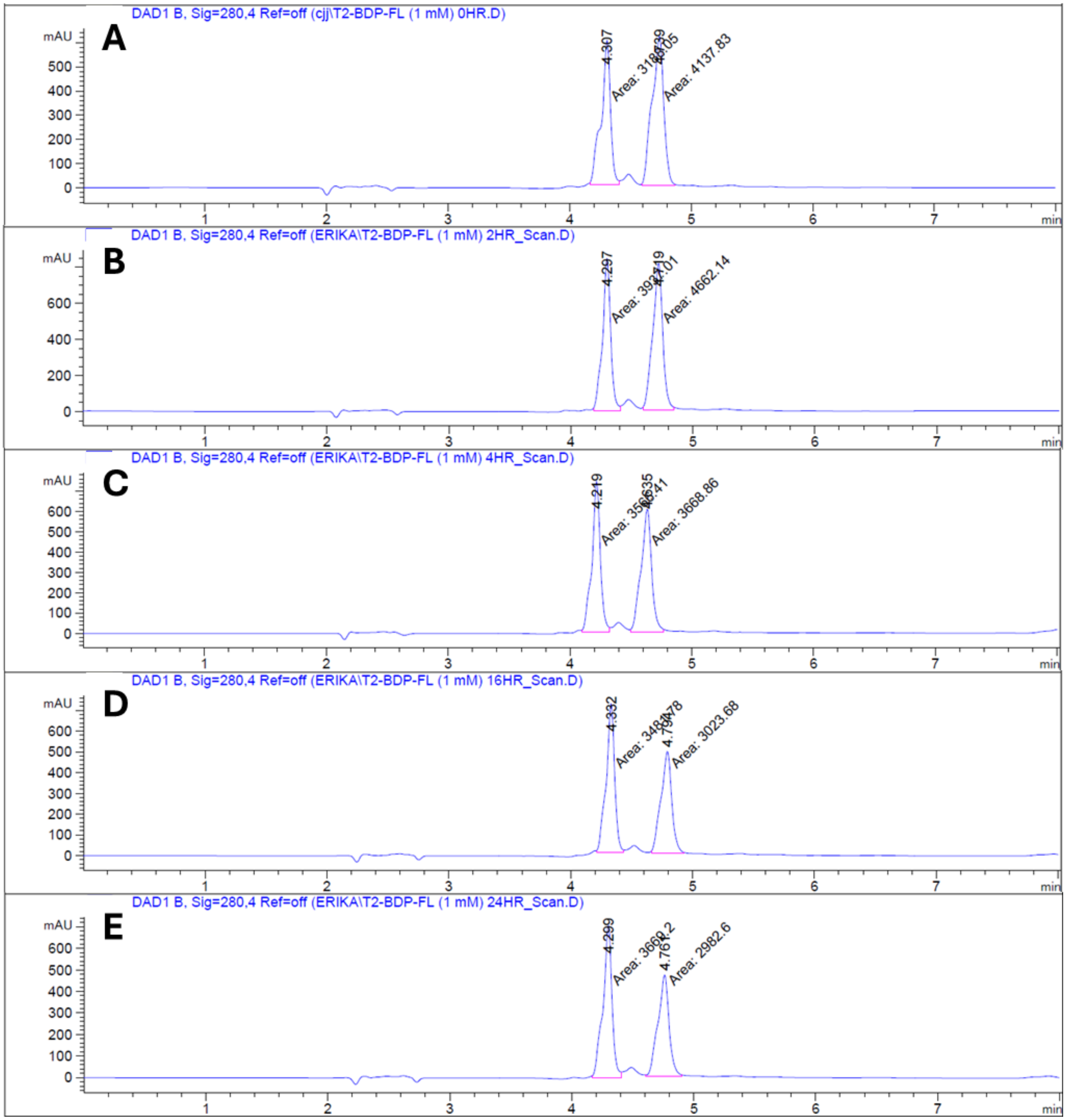

[a] LC–UV chromatograms showing the stability of the T2-BDP-FL (1 mM) tracer in cell medium over time (panels A–E: 0, 2, 4, 16, and 24 h, respectively). Imipramine was used as an internal standard. The T2-BDP-FL tracer eluted at approximately 4.7 min, and Imipramine at approximately 4.3 min. A time-dependent decrease in tracer peak area relative to the internal standard indicates degradation over the 24-hour period.

*Figure S6. Time-Dependent Degradation of T2-BDP-589 Tracer in Cell Medium Monitored by LC-UV [a]*

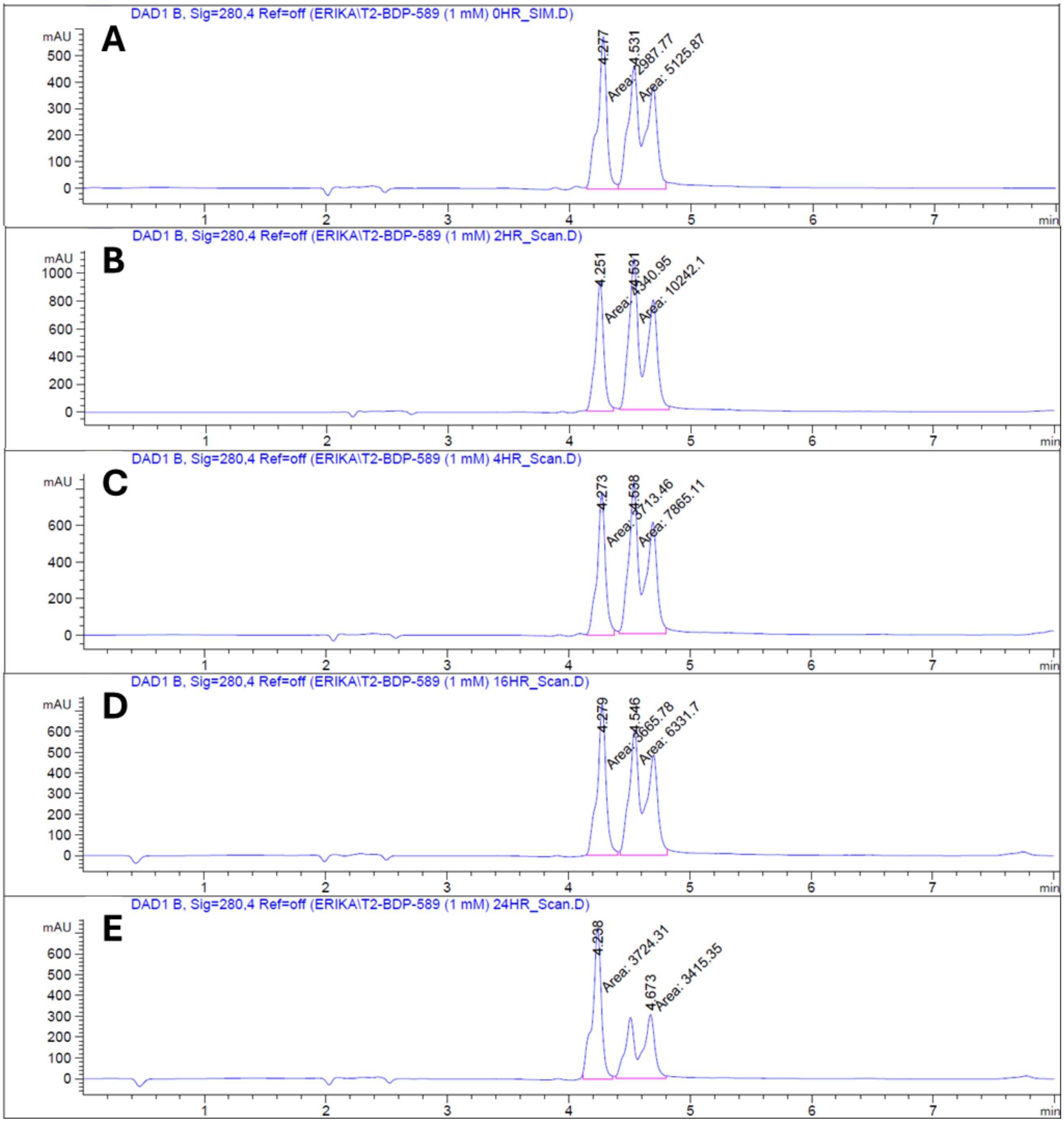

[a] LC–UV chromatograms showing the stability of the T2-BDP-589 (1 mM) tracer in cell medium over time (panels A–E: 0, 2, 4, 16, and 24 h, respectively). Imipramine was used as an internal standard. The T2-BDP-589 tracer eluted at approximately 4.5 min, and Imipramine at approximately 4.3 min. A time-dependent decrease in tracer peak area relative to the internal standard indicates degradation over the 24-hour period.

*Figure S7. Time-Dependent Degradation of K5-BDP-FL Tracer in Cell Medium Monitored by LC-UV [a]*

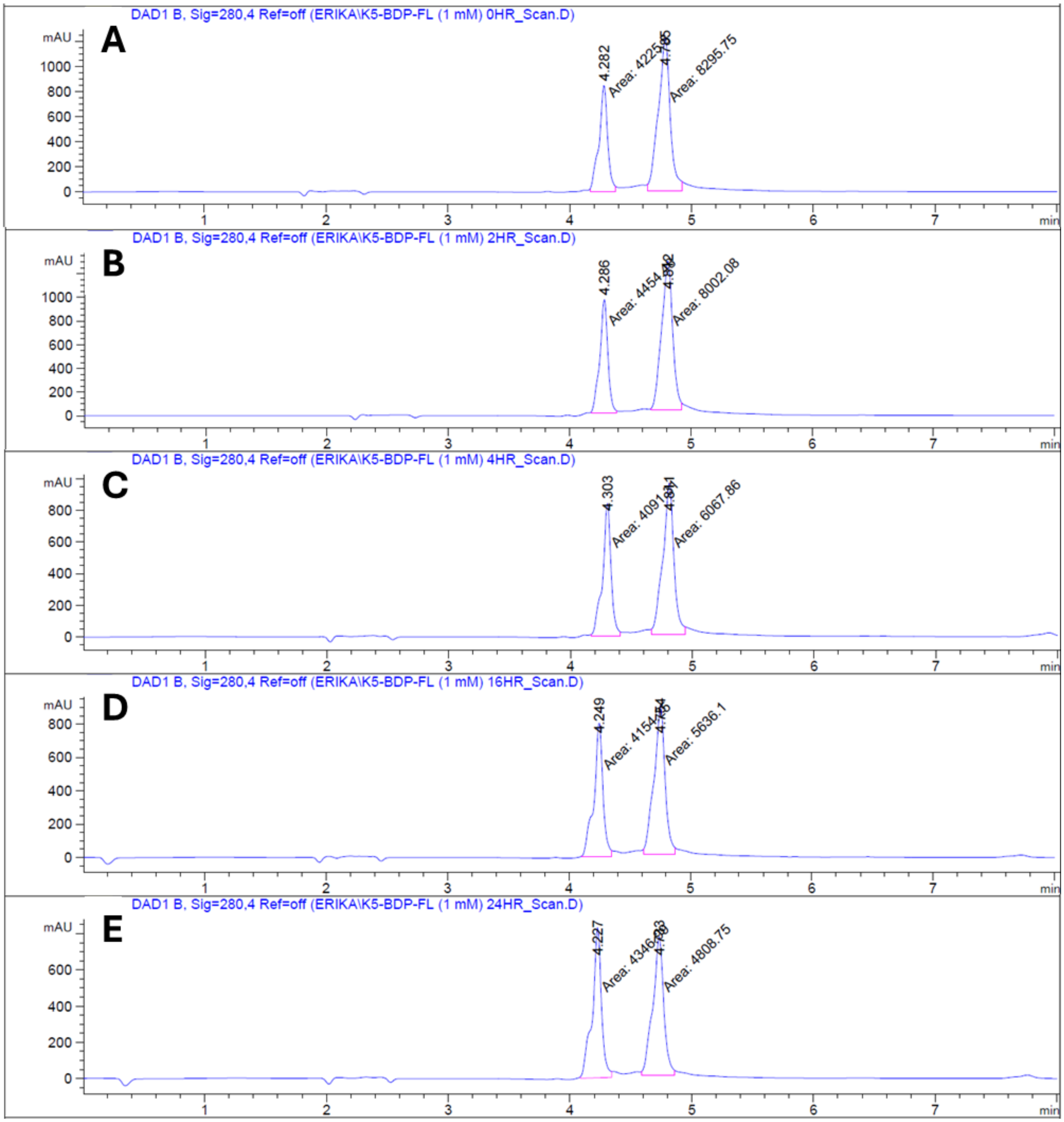

[a] LC–UV chromatograms showing the stability of the K5-BDP-FL (1 mM) tracer in cell medium over time (panels A–E: 0, 2, 4, 16, and 24 h, respectively). Imipramine was used as an internal standard. The K5-BDP-FL tracer eluted at approximately 4.8 min, and Imipramine at approximately 4.3 min. A time-dependent decrease in tracer peak area relative to the internal standard indicates degradation over the 24-hour period.

*Figure S8. Time-Dependent Degradation of K5-BDP-589 Tracer in Cell Medium Monitored by LC-UV [a]*

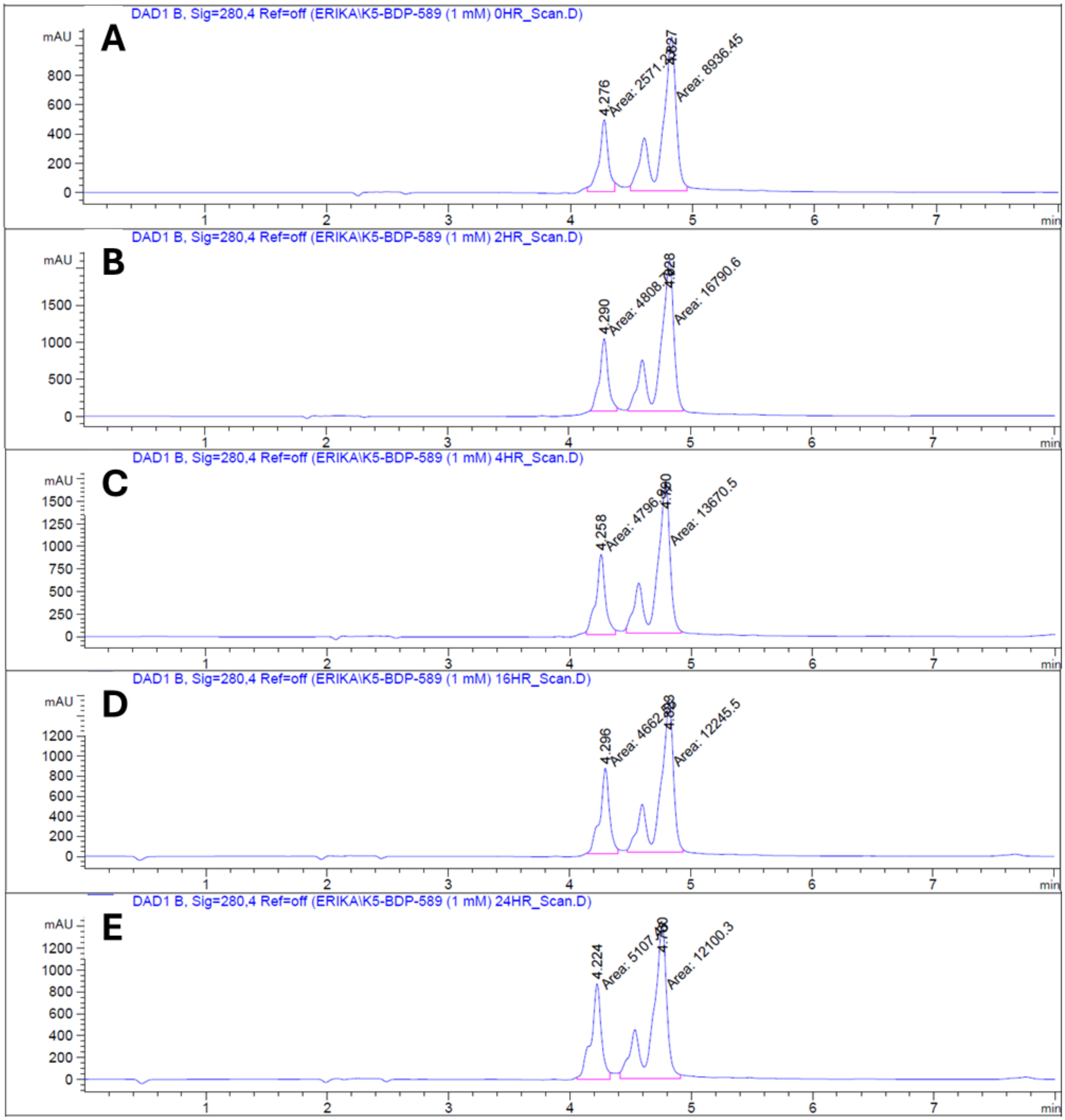

[a] LC–UV chromatograms showing the stability of the K5-BDP-589 (1 mM) tracer in cell medium over time (panels A–E: 0, 2, 4, 16, and 24 h, respectively). Imipramine was used as an internal standard. The K5-BDP-589 tracer eluted at approximately 4.8 min, and Imipramine at approximately 4.3 min. A time-dependent decrease in tracer peak area relative to the internal standard indicates degradation over the 24-hour period.

***Table S1. Criteria for Signal Window and Z'*** [a]

| *Signal Window* | *Z'* |
|---|---|
| Recommended: SW > 2 | Excellent: Z' > 0.5 |
| Acceptable: SW > 1 | Do-able: 0 < Z' < 0.5 |
| Unacceptable: SW < 1 | Unacceptable: Z' < 0 |

SW = signal window

[a] The Signal Window is classified into three categories: Recommended, which ensures optimal assay performance; Acceptable, which meets minimum quality requirements; and Unacceptable, which indicates poor assay conditions requiring optimization. The Z' factor is categorized as Excellent (> 0.5), indicating robust assay performance; Do-able (0 – 0.5), signifying moderate quality with potential for improvement; and Unacceptable (< 0), representing inadequate assay performance.

***Table S2. Z' for RIPK1 TR-FRET Assay [a]***

| *Probes* | *Filter Settings* | *Maximum Signal* | | *Background Signal* | | | |
|---|---|---|---|---|---|---|---|
| | | *Mean* | *SD* | *Mean* | *SD* | *Z'* | *SW* |
| T2-BODIPY-FL | 520 ± 10 nm/490 ± 10 nm | 9.01x $10^4$ | 3930 | 4.89 X $10^4$ | 1920 | 0.57 | 6.0 |
| | 640 ± 10 nm/490 ± 10 nm | 2.00 X $10^4$ | 417 | 1.94 X $10^4$ | 142 | -1.7 | -2.5 |
| | 640 ± 10 nm/550 ± 25 nm | 3.80 X $10^4$ | 929 | 3.97 X $10^4$ | 302 | -1.2 | -2.2 |
| T2-BODIPY-589 | 520 ± 10 nm/490 ± 10 nm | 1.80 X $10^4$ | 141 | 2.17 X $10^4$ | 589 | 0.40 | 10.5 |
| | 640 ± 10 nm/490 ± 10 nm | 5.46 X $10^4$ | 1030 | 3.94 X $10^4$ | 1340 | 0.53 | 7.8 |
| | 640 ± 10 nm/550 ± 25 nm | 1.22 X $10^5$ | 2750 | 9.35 X $10^4$ | 3950 | 0.29 | 3.1 |

[a] The optimized results in the Time-Resolved Fluorescent Resonance Energy Transfer (TR-FRET) assay were conducted at a probe concentration of 2 μM. The table summarizes the mean and standard deviation (SD) of the maximum signal and background for each fluorescent tracer, alongside the calculated Z' factor and signal window for various filter settings.

***Table S3. Optimized TR-FRET Assay Reveals Stronger RIPK1 Unlabeled T2 Inhibitor Binding with T2-BODIPY-FL Over T2-BODIPY-589***

| *Probes* | *Filter Settings* | *T2 Inhibitor Ki (nM)* |
|---|---|---|
| T2-BODIPY-FL | 520 ± 10 nm/490 ± 10 nm | 18 ± 0.70 |
| T2-BODIPY-589 | 640 ± 10 nm/490 ± 10 nm | 54 ± 5.9 |

***Table S4. Z' for BTK TR-FRET Assay [a]***

| | | *Maximum Signal* | | *Background Signal* | | | |
|---|---|---|---|---|---|---|---|
| *Probes* | *Filter Settings* | *Mean* | *SD* | *Mean* | *SD* | *Z'* | *SW* |
| K5-BODIPY-FL | 520 ± 10 nm/490 ± 10 nm | $6.44 \times 10^4$ | 1380 | $2.36 \times 10^4$ | 439 | 0.87 | 25.52 |
| | 640 ± 10 nm/490 ± 10 nm | $1.84 \times 10^4$ | 332 | $1.80 \times 10^4$ | 122 | -2.03 | N/A |
| | 640 ± 10 nm/550 ± 25 nm | $3.56 \times 10^4$ | 710 | $3.65 \times 10^4$ | 286 | -2.17 | N/A |
| K5-BODIPY-589 | 520 ± 10 nm/490 ± 10 nm | $1.50 \times 10^4$ | 454 | $1.78 \times 10^4$ | 640 | -0.16 | N/A |
| | 640 ± 10 nm/490 ± 10 nm | $2.84 \times 10^4$ | 401 | $2.12 \times 10^4$ | 441 | 0.65 | 11.51 |
| | 640 ± 10 nm/550 ± 25 nm | $5.62 \times 10^4$ | 934 | $4.41 \times 10^4$ | 652 | 0.61 | 7.88 |

[a] The optimized results in the Time-Resolved Fluorescent Resonance Energy Transfer (TR-FRET) assay were conducted at a probe concentration of 0.125 μM. The table summarizes the mean and standard deviation (SD) of the maximum signal and background for each fluorescent tracer, alongside the calculated Z' factor and signal window for various filter settings.

***Table S5. Optimized TR-FRET Assay Reveals Stronger BTK Ibrutinib Inhibitor Binding with K5-BODIPY-FL Over K5-BODIPY-589***

| *Probes* | *Filter Settings* | *Ibrutinib Ki (nM)* |
|---|---|---|
| K5-BODIPY-FL | 520 ± 10 nm/490 ± 10 nm | 30.07 ± 1.98 |
| K5-BODIPY-589 | 640 ± 10 nm/490 ± 10 nm | 84.05 ± 8.44 |
| K5-BODIPY-589 | 640 ± 10 nm/550 ± 25 nm | 60.38 ± 5.68 |

***Table S6. Z' for HEK293 nLuc-RIPK1 NanoBRET Assay [a]***

| *Probes* | *Monochromatic Settings with Selected Filter* | *Maximum Signal* | | *Background Signal* | | | |
|---|---|---|---|---|---|---|---|
| | | *Mean* | *SD* | *Mean* | *SD* | *Z'* | *SW* |
| T2-BODIPY-FL | 520 ± 30 nm/460 ± 30 nm | 238 | 12.4 | -8.95 | 10.3 | 0.72 | 14 |
| | 550 ± 30 nm/460 ± 30 nm | 72.0 | 2.85 | -0.40 | 5.24 | 0.66 | 17 |
| | 580 ± 30 nm/460 ± 30 nm | 15.5 | 0.60 | -0.30 | 1.28 | 0.64 | 17 |
| | 610 nm LP/460 ± 30 nm | 13.6 | 1.00 | -1.11 | 2.87 | 0.21 | 3.1 |
| T2-BODIPY-589 | 580 ± 30 nm/460 ± 30 nm | 15.3 | 1.32 | -0.78 | 0.73 | 0.62 | 7.6 |
| | 610 ± 30 nm/460 ± 30 nm | 21.2 | 0.94 | 1.26 | 0.46 | 0.79 | 17 |
| | 610 nm LP/460 ± 30 nm | 151 | 8.09 | 1.44 | 1.89 | 0.80 | 15 |
| | 640 ± 30 nm/460 ± 30 nm | 5.42 | 0.38 | 0.36 | 0.32 | 0.58 | 7.7 |

[a] The optimization results of the Nano Bioluminescent Resonance Energy Transfer (NanoBRET) assay conducted at a probe concentration of 1 µM. The table presents the mean and standard deviation of the maximum signal and background for each fluorescent tracer, along with the calculated Z' factor and signal window (SD) for various monochromatic conditions. A 610 nm long-pass filter was employed to optimize a single condition for each tracer.

***Table S7. Optimized NanoBRET Assay Confirms Consistent RIPK1 Unlabeled T2 Inhibitor Binding Across Probes***

| *Probes* | *Monochromatic Settings with Selected Filter* | *T2 Inhibitor Ki (nM)* |
|---|---|---|
| T2-BODIPY-FL | 520 ± 30 nm/460 ± 30 nm | 14 ± 2.5 |
| T2-BODIPY-FL | 550 ± 30 nm/460 ± 30 nm | 13 ± 3.4 |
| T2-BODIPY-FL | 580 ± 30 nm/460 ± 30 nm | 9.1 ± 4.2 |
| T2-BODIPY-589 | 610 ± 30 nm/460 ± 30 nm | 6.6 ± 1.6 |
| T2-BODIPY-589 | 610 nm LP/460 ± 30 nm | 10 ± 1.5 |
| T2-BODIPY-589 | 640 ± 30 nm/460 ± 30 nm | 11 ± 2.8 |

***Table S8. Z' for HeLa BTK-nLuc NanoBRET Assay*** [a]

| *Probes* | *Monochromatic Settings with Selected Filter* | *Maximum Signal* | | *Background Signal* | | | |
|---|---|---|---|---|---|---|---|
| | | *Mean* | *SD* | *Mean* | *SD* | *Z'* | *SW* |
| K5-BODIPY-FL | 520 ± 20 nm/460 ± 30 nm | 56.60 | 5.33 | 0.19 | 0.93 | 0.67 | 7.06 |
| | 540 ± 20 nm/460 ± 30 nm | 26.98 | 2.21 | 2.17 | 0.86 | 0.63 | 7.08 |
| | 560 ± 20 nm/460 ± 30 nm | 10.69 | 1.41 | 0.63 | 1.04 | 0.27 | 1.92 |
| | 580 ± 20 nm/460 ± 30 nm | 3.66 | 0.87 | 0.22 | 0.74 | -0.41 | -1.60 |
| K5-BODIPY-589 | 580 ± 30 nm/460 ± 30 nm | 14.79 | 1.11 | -0.39 | 1.20 | 0.54 | 7.46 |
| | 610 ± 30 nm/460 ± 30 nm | 22.43 | 1.21 | 0.28 | 0.27 | 0.80 | 14.61 |
| | 610 nm LP/460 ± 30 nm | 145.37 | 6.90 | 3.67 | 1.32 | 0.83 | 16.97 |
| | 640 ± 30 nm/460 ± 30 nm | 5.84 | 1.04 | 0.31 | 0.11 | 0.38 | 2.01 |

[a] The optimization results of the Nano Bioluminescent Resonance Energy Transfer (NanoBRET) assay conducted at a probe concentration of 1 μM. The table presents the mean and standard deviation of the maximum signal and background for each fluorescent tracer, along with the calculated Z' factor and signal window (SD) for various filter settings.

***Table S9. Optimized NanoBRET Assay Confirms Consistent BTK Ibrutinib Inhibitor Binding Across Probes***

| *Probes* | *Monochromatic Settings with Selected Filter* | *Ibrutinib Ki (nM)* |
|---|---|---|
| K5-BODIPY-FL | 520 ± 20 nm/460 ± 30 nm | 0.40 ± 0.05 |
| K5-BODIPY-FL | 540 ± 20 nm/460 ± 30 nm | 0.53 ± 0.10 |
| K5-BODIPY-FL | 560 ± 20 nm/460 ± 30 nm | 0.27 ± 0.06 |
| K5-BODIPY-589 | 580 ± 30 nm/460 ± 30 nm | 0.50 ± 0.12 |
| K5-BODIPY-589 | 610 ± 30 nm/460 ± 30 nm | 0.29 ± 0.04 |
| K5-BODIPY-589 | 610 nm LP/460 ± 30 nm | 0.40 ± 0.05 |
| K5-BODIPY-589 | 640 ± 30 nm/460 ± 30 nm | 0.47 ± 0.11 |

***Table S10: LC-MS Quantification of Probe Concentration Over Time Using Imipramine as an Internal Standard***

| *Probe* | *Time (H)* | *% Degradation* |
|---|---|---|
| T2-BDP-FL | 0 | 0.00 |
| | 2 | 8.76 |
| | 4 | 20.74 |
| | 16 | 33.09 |
| | 24 | 37.37 |
| T2-BDP-589 | 0 | 0.00 |
| | 2 | 0.00 |
| | 4 | 10.23 |
| | 16 | 26.79 |
| | 24 | 61.13 |
| K5-BDP-FL | 0 | 0.00 |
| | 2 | 8.50 |
| | 4 | 24.46 |
| | 16 | 30.90 |
| | 24 | 43.65 |
| K5-BDP-589 | 0 | 0.00 |
| | 2 | 0.00 |
| | 4 | 18.00 |
| | 16 | 24.43 |
| | 24 | 31.83 |

### EXPERIMENTAL METHODS

#### *ClarioSTAR Plus Plate Reader from BMG LABTECH*

The ClarioSTAR Plus plate reader from BMG LABTECH is a sophisticated multimode instrument tailored for diverse applications in biochemical and cellular assays. It features a high-performance monochromator capable of exciting and detecting wavelengths ranging from 220 nm to 1,000 nm, with adjustable bandwidth settings from 5 nm to 100 nm. The plate reader supports multiple detection modes, including fluorescence, luminescence, absorbance, and time-resolved fluorescence, enabling the execution of assays such as Time-Resolved Fluorescence Resonance Energy Transfer (TR-FRET) and Nano Bioluminescent Resonance Energy Transfer (NanoBRET). Its advanced optical system combines filters and a monochromator, allowing for optimal assay condition adjustments and ensuring accurate data acquisition. Additionally, the ClarioSTAR Plus offers high-resolution top- and bottom-reading capabilities across various plate formats (96-, 384-, and 1,536-well plates). With an intuitive interface and automated features, it enhances data acquisition and analysis, making it an essential tool for high-throughput screening and kinetic studies in drug discovery.

#### *Optical Filters*

To optimize the detection parameters for TR-FRET, optical filters were selected based on their spectral properties, optical density, and compatibility with the fluorescence detection system. The optical filters used in this study were procured from two manufacturers: BMG Labtech and Avantier Inc.

**Filters from BMG LABTECH.** The excitation and emission filters from BMG Labtech were chosen for their high-transmission efficiency and precision optical coatings. The specific filters used include:

- **Excitation Filter (EX TR):** Bandpass filter (CWL = 330 nm, FWHM = 75 nm), offering high-performance excitation for time-resolved fluorescence (TRF) assays.
- **Emission Filter (490 ± 10 nm):** Bandpass filter (CWL = 490 nm, FWHM = 10 nm), offering high optical density for improved signal-to-noise ratio.
- **Emission Filter (520 ± 10 nm):** Bandpass filter (CWL = 520 nm, FWHM = 10 nm), provides high sensitivity for detecting weak fluorescence signals.
- **Emission Filter (610 nm LP):** A long-pass filter with a cut-on wavelength of 610 nm, designed to facilitate the detection of red and far-red fluorescent acceptors in NanoBRET assays by effectively capturing acceptor emissions while minimizing background interference.
- **Dichroic Mirror (LP TR):** 426 nm long-pass filter to effectively separate excitation and emission light paths.

**Filters from Avantier Inc.** Avantier Inc. provided an alternative set of filters with similar specifications but different coating technologies to enhance blocking and transmission. The filters used include:

- **Emission Filter (550 ± 25 nm):** Bandpass filter (CWL = 550 nm, FWHM = 25±4 nm), optimized for emission detection while minimizing background interference.
- **Emission Filter (640 ± 10 nm):** Bandpass filter (CWL = 640 nm, FWHM = 10±2 nm), designed for efficient excitation of T2-BDP-589.

#### *Monochromator*

The ClarioSTAR Plus features a Linear Variable Filter (LVF) monochromator, which is a key component of its advanced optical system and allows for excitation and emission wavelength selection across a broad range. The monochromator settings utilized in this study included:

- **460 ± 30 nm:** Employed for NanoLuc luminescence detection in NanoBRET assays, providing optimal excitation for the NanoLuc donor.
- **Variable Bandwidth:** The monochromator supports variable bandwidth settings, which can be adjusted incrementally (e.g., 30 nm intervals) to fine-tune detection parameters. This flexibility allows for precise adjustments to minimize bleed-through and optimize signal-to-noise ratios during assays.

*Materials*

| Reagent or Resource | Source | Identifier |
|---|---|---|
| **Antibodies** | | |
| HTRF mAb Anti-6His Tb-Conjugate, 1,000 Assay Points | Revvity | Cat. # 61HISTLF |
| HTRF Anti-GST mAb Tb-Conjugate, 1,000 Assay Points | Revvity | Cat. # 61GSTTLF |
| **Cell line** | | |
| HEK 293T | ATCC | Cat. # CRL-3216 |
| HeLa | ATCC | Cat. # CCL-2 |
| **Reagents, proteins, and medium** | | |
| DMEM | Corning | Cat. # 10-013-CV |
| Fetal Bovine Serum (FBS) | Corning | Cat. # 35-011-CV |
| Gateway$^{TM}$ LR Clonase$^{TM}$ II Enzyme mix | ThermoFisher | Cat. # 11791020 |
| Lipomaster 3000 Transfection Reagent | Vazyme | Cat. # TL301-02 |
| Opti-MEM | ThermoFisher | Cat. # 11058021 |
| pLeni6.2-ccdB-nLuc plasmid | Addgene | Cat. # 87075 |
| Recombinant 6His-tagged human RIPK1 protein | In house | |
| GST-TEV-GS-BTK protein | In house | |
| T2-BDP-FL | In house | |
| T2-BDP-589 | In house | |
| K5-BDP-FL | In house | |
| K5-BDP-576/589 | Promega | Cat. # N2482 |
| 0.25% Trypsin, 2.21 mM EDTA, 1X [-] sodium bicarbonate | Corning | Cat. # 25-053-CI |
| Unlabeled T2 Inhibitor | In house | |
| Ibrutinib | In house | |
| NanoBRET® Nano-Glo® Substrate | Promega | Cat. # N1571 |
| NanoLuc® extracellular inhibitor | Promega | Cat. # N2162 |
| **Plasmids** | | |
| nLuc/hRIPK1 plasmid | Promega | Cat. # NV4171 |
| BTK/nLuc plasmid | Viva Biotech | 241015 |
| **Instrumentation and software** | | |
| CLARIOstar *Plus* | BMG LABTECH | Cary, NC |
| GraphPad Prism 10 | GraphPad Software | San Diego, CA |

***Reagents***

Unless stated overwise, all chemicals were sourced from Sigma-Aldrich. Terbium (Tb) cryptate-labeled anti-6His-antibody (anti-His-Tb) was purchased from Revvity. T2 tracers and unlabeled T2 inhibitor were synthesized, as previously described.[1] K5 tracers were synthesized and purchased from Promega.

***Cell culture and medium***

HEK293T cells (ATCC, Cat. No. CRL-11268) and HeLa cells (ATCC, Cat. No. CCL-2) were cultured in Dulbecco's Modified Eagle Medium (DMEM) (Corning, Cat. No. 10-013-CV) supplemented with 10% fetal bovine serum (FBS) (Corning, Cat. No. 35-011-CV) and phenol red, at 37°C with 5% $CO_2$. HEK293T cells require periodic passaging, usually every 2-3 days, when they reach 70-80% confluency. During passaging, the cells are gently detached using trypsin-EDTA (0.25%) solution (Corning, Cat. No. 25-053-CI), neutralized with medium containing 10% FBS, and then seeded at an appropriate density for continued culture or experimental use. The medium is replaced every 2-3 days to ensure optimal growth conditions.

***Expression and Purification of His-hRIPK1 and GST-BTK Proteins***

Recombinant 6His-tagged hRIPK1 was expressed and purified essentially as previously described.[1]

The GST-tagged BTK protein was produced by Viva Biotech CRO protein expression and purification services. The purified BTK protein final concentration was 0.07 mg/mL and had the following amino acid sequence:
MSPILGYWKIKGLVQPTRLLLEYLEEKYEEHLYERDEGDKWRNKKFELGLEFPNLPYYIDGDVKLTQSMAIIRYIA
DKHNMLGGCPKERAEISMLEGAVLDIRYGVSRIAYSKDFETLKVDFLSKLPEMLKMFEDRLCHKTYLNGDHVTH
PDFMLYDALDVVLYMDPMCLDAFPKLVCFKKRIEAIPQIDKYLKSSKYIAWPLQGWQATFGGGDHPPKSDENLYF
QGGSMAAVILESIFLKRSQQKKKTSPLNFKKRLFLLTVHKLSYYEYDFERGRRGSKKGSIDVEKITCVETVVPEKN
PPPERQIPRRGEESSEMEQISIIERFPYPFQVVYDEGPLYVFSPTEELRKRWIHQLKNVIRYNSDLVQKYHPCFWIDG
QYLCCSQTAKNAMGCQILENRNGSLKPGSSHRKTKKPLPPTPEEDQILKKPLPPEPAAAPVSTSELKKVVALYDY
MPMNANDLQLRKGDEYFILEESNLPWWRARDKNGQEGYIPSNYVTEAEDSIEMYEWYSKHMTRSQAEQLLKQ
EGKEGGFIVRDSSKAGKYTVSVFAKSTGDPQGVIRHYVVCSTPQSQYYLAEKHLFSTIPELINYHQHNSAGLISRL
KYPVSQQNKNAPSTAGLGYGSWEIDPKDLTFLKELGTGQFGVVKYGKWRGQYDVAIKMIKEGSMSEDEFIEEAK
VMMNLSHEKLVQLYGVCTKQRPIFIITEYMANGCLLNYLREMRHRFQTQQLLEMCKDVCEAMEYLESKQFLHR
DLAARNCLVNDQGVVKVSDFGLSRYVLDDEYTSSVGSKFPVRWSPPEVLMYSKFSSKSDIWAFGVLMWEIYSLG
KMPYERFTNSETAEHIAQGLRLYRPHLASEKVYTIMYSCWHEKADERPTFKILLSNILDVMDEES

***Mammalian protein expression***

The nLuc/hRIPK1 (Promega, Cat. No. NV4171) and BTK/nLuc (Promega, Cat. No. N2441) constructs were used for transfection experiments. The nLuc/hRIPK1 and BTK/nLuc constructs was generated by cloning the human RIPK1 cDNA insert or human BTK cDNA insert into the pLeni6.2-ccdB-nLuc plasmid (Addgene #87075) using the Gateway Cloning Kit (Thermo #11791020) in a solution of 0.125 M $CaCl_2$ and 1x HBSS. For a 6 cm culture dish, 2 million HEK293T/HeLa cells were seeded in 5 mL of DMEM the day prior to transfection. Transfections were conducted when the cells reached 70%-90% confluence. Transfection was performed using the Lipomaster 3000 protocol (Vazyme, Cat. No. TL301-02. To prepare the transfection mixture, 8.25 µL of Lipomaster 3000 reagent was gently mixed with 250 µL of Opti-MEM (ThermoFisher, Cat. No. 11058021), followed by the addition of the appropriate amount of plasmid DNA combined with 11 µL of T3000 Enhancer reagent and an additional 250 µL of Opti-MEM. The Lipomaster 3000 reagent/DNA/T3000 enhancer complex was incubated at room temperature for 15 minutes. The resulting transfection cocktail was then added dropwise to the culture dish and evenly distributed by gently shaking. Following this, the cells were incubated for 48 hours to allow for gene expression before proceeding with downstream assays.

***Optimization of TR-FRET Filter Pairs for RIPK1 Detection Using Fluorescent Tracers***

The TR-FRET assay was performed using 384-well white, flat bottom, non-binding surface plates (Corning, Cat. No. 3824) with a total reaction volume of 20 µL in each well, six measurements. To optimize TR-FRET filter pairs for protein detection using fluorescent tracers, 1 nM of His-hRIPK protein was prepared in reaction buffer (50 mM Tris, pH 7.5, 0.1% Trition X-100, 0.01% BSA, and 1 mM TCEP). The anti-His-Tb antibody was diluted to a final concentration of 0.33 nM in the same buffer and incubated with the His-hRIPK1 protein at 0° C. A 5 µL aliquot of the protein/antibody mixture was then added to designated wells. To assess the binding affinity of hRIPK1 to T2-BDP-FL and T2-BDP-589, both tracers were serially diluted in reaction buffer (final concentrations ranging from 4 µM to 0.00039 µM, 2-fold, 11 pts) with 0.1% DMSO, and 5 µL of each tracer, at appropriate concentrations, were added to each well. To eliminate background off-target binding, 5 µL of unlabeled T2 inhibitor (final concentration: 10 uM) was added to the designated wells, while 5 µL of reaction buffer was added to the remaining wells. The fluorescence intensity of the donor and acceptors was measured every hour over a 4-hour period, at room temperature, using a CLARIOstar *Plus* microplate reader (BMG LABTECH, Cary, NC) equipped with TR-FRET detection capability: EX TR excitation, 490±10 nm (Tb) or 550±25 nm (Tb) and either 520±10 nm (T2-BDP-FL) or 640±10 nm (T2-BDP-589) emission, delay 100 µs, integration 400 µs. The TR-FRET signal was calculated by recording both donor and acceptor emissions, and the TR-FRET ratio (acceptor emission/donor emission) was analyzed using a One site- Total and Nonspecific Binding equation in GraphPad Prism 10 (GraphPad Software, San Diego, CA) to determine the dissociation constant ($K_d$). The filter pair that yielded the highest Z' factor was selected as the optimal configuration.

***TR-FRET Competition Assay for RIPK1 Using Optimized Filter Settings for T2-BDP-FL and T2-BDP-589***

A TR-FRET competition assay was conducted to assess the binding interactions between the target protein and the unlabeled T2 inhibitor. Initially, 1 nM of His-hRIPK1 protein was incubated with 0.33 nM of anti-His-Tb antibody at 0°C. A 5 µL aliquot of the protein/antibody mixture was subsequently added to the designated well of a 384-well white, flat bottom, non-binding surface plate (Corning, Cat. No. 3824, assay volume of 20 µL, six measurements). To each well, 5 µL of a fixed $K_d$ concentration of T2-BDP-FL or T2-BDP-589 tracers in reaction buffer (50 mM Tris, pH 7.5, 0.1% Trition X-100, 0.01% BSA, and 1 mM TCEP) was introduced. Following this, 5 µL of unlabeled T2 inhibitor and DMSO were added at varying concentrations (ranging from 10 µM to 0.0097 µM, 2-fold, 11 points). Fluorescence intensities of both donor and acceptor were measured using a CLARIOstar Plus microplate reader (BMG LABTECH, Cary, NC), with emissions recorded at EX TR excitation, 490±10 nm (Tb) and either 520±10 nm (T2-BDP-FL) or 640±10 nm (T2-BDP-589) emission, delay 100 µs, integration 400 µs. The TR-FRET ratio (10,000 × acceptor emission/donor emission) was calculated every hour over a 4-hour period. The competition between the inhibitor and the fluorescent tracer was evaluated by monitoring the decrease in the TR-FRET signal as the inhibitor concentration increased, and data were normalized to the negative control (DMSO) to determine the percent inhibition at each inhibitor concentration. Normalized percent inhibition was plotted against inhibitor concentration and fitted to a sigmoidal dose-response curve [inhibitor] vs. normalized response equation. To accurately calculate the inhibitory constant ($K_i$), fluorescent tracer concentrations corresponding to their dissociation constants ($K_d$) were selected, and the Cheng-Prusoff equation ($Ki = IC_{50}/2$) was applied.[2] This approach facilitated the identification and quantification of inhibitors based on their ability to compete with the labeled tracer for binding to the target protein.

To further demonstrate the generalizability of these detection parameters beyond RIPK1, we extended TR-FRET assay development to Bruton's tyrosine kinase (BTK). Using GST-tagged BTK and the established BTK tracers K5-BODIPY-FL (K5-BDP-FL) and K5-BODIPY-576/589 (K5-BDP-589), we evaluated assay performance under similar TR-FRET conditions.[3] Data in Figures S1-S2 and Tables S4-S5 revealed comparable binding characteristics and robust signal windows, validating both tracers' effectiveness and filter configurations for BTK. These findings reinforced that the optimized TR-FRET assay conditions are suitable for RIPK1 and reliably applicable to additional kinase targets.

***Optimizing NanoBRET Assay for Accurate RIPK1 Target Engagement with Fluorescent Tracers***

HEK293T cells (ATCC, Cat. No. CRL-3216) were transiently transfected with the nLuc/hRIPK1 plasmid according to the mammalian protein expression procedure. After incubation, the medium was aspirated, and the cells were trypsinized with a trypsin-EDTA (0.25%) solution (Corning, Cat. No. 25-053-CI). The cells were then resuspended in Opti-MEM medium (ThermoFisher, Cat. No. 11058021) at a density of $2 \times 10^5$ cells/mL. A volume of 80 µL of the cell suspension was plated into 96-well plates (Corning, Cat. No. 3600, total assay volume of 100 µL, four replicates). Serially diluted T2-BDP-FL and T2-BDP-589 tracers (final concentrations ranging from 5 µM to 0.0000026 µM, 5-fold, 10 points) and 0.1% DMSO were prepared. A 10 µL aliquot of each tracer, at the appropriate concentration, was added to the corresponding wells. To minimize background off-target binding, 10 µL of unlabeled T2 inhibitor (final concentration: 10 µM) was added to the designated wells, while 10 µL of Opti-MEM medium (ThermoFisher, Cat. No. 11058021) was added to the remaining wells. The plate was incubated for 2 hours at 37°C with 5% $CO_2$.

After incubation, BRET measurements were performed by adding NanoBRET® NanoGlo® Substrate (Promega, Cat. No. N1571) and NanoLuc® extracellular inhibitor (Promega, Cat. No. N2162). Luminescence and fluorescence intensity were measured using a CLARIOstar Plus microplate reader (BMG LABTECH, Cary, NC) equipped with LVF monochromator detection capabilities: 460 ± 30 nm (nLuc donor), 520 ± 30 nm, 550 ± 30 nm, 580 ± 30 nm, 610 ± 30 nm, 610 nm LP, and 640 ± 30 nm emission.

MilliBRET units (mBU) were calculated by multiplying the BRET values by 1,000. The resulting mBU signals were analyzed using the One-site Total Binding equation in GraphPad Prism 10 software (GraphPad Software, San Diego, CA) to determine the dissociation constant (Kd). This analysis generated binding curves for each experimental group and enabled the calculation of the corresponding dissociation constants ($K_d$ values). The filter pair that yielded the highest Z' factor was selected as the optimal configuration.

***NanoBRET assay for RIPK1 with Optimized Filter Settings***

To assess the binding interactions between nLuc/hRIPK1 and the unlabeled T2 inhibitor, 80 µL of HEK293T cells expressing the nLuc/hRIPK1 target protein are seeded into 96-well plates (Corning, Cat. No. 3600, total assay volume of 100 µL, four replicates) at a density of $2 \times 10^5$ cells/mL in Opti-MEM medium (ThermoFisher, Cat. No. 11058021). To each well, 10 µL of a fixed $K_d$ concentration of T2-BDP-FL and T2-BDP-589 tracers in Opti-MEM medium (ThermoFisher, Cat. No. 11058021) was added. Subsequently, 10 µL of serial dilutions of unlabeled T2 inhibitor and DMSO were added at varying concentrations (ranging from 10 µL to $5.1 \times 10^{-7}$ µM, 5-fold, 10 points). The plate was incubated for 2 hours at 37°C with 5% $CO_2$.

Following incubation, BRET measurements were conducted by adding NanoBRET® NanoGlo® Substrate (Promega, Cat. No. N1571) and NanoLuc® extracellular inhibitor (Promega, Cat. No. N2162). Luminescence and fluorescence intensity were measured using a CLARIOstar Plus microplate reader (BMG LABTECH, Cary, NC) equipped with LVF monochromator detection capabilities: 460±30 nm (nLuc donor), 520±30 nm, 550±30 nm, 580±30 nm, 610±30 nm, LP610, and 640±30 nm emission.

The competition between the inhibitor and the fluorescent tracer was evaluated by monitoring the decrease in the BRET signal as the inhibitor concentration increased. Data were normalized to the negative control (DMSO) to determine the percent inhibition at each inhibitor concentration. Normalized percent inhibition was plotted against inhibitor concentration and fitted to a sigmoidal dose-response curve ([inhibitor] vs. normalized response). To accurately calculate the inhibitory constant ($K_i$), fluorescent tracer concentrations corresponding to their dissociation constants ($K_d$) were selected, and the Cheng-Prusoff equation ($Ki = IC_{50}/2$) was applied.[2] This method allowed for the identification and quantification of inhibitors based on their ability to compete with the labeled tracer for binding to the target protein.

To further validate the cross-platform applicability and broader utility, we extended NanoBRET target engagement assays into HeLa cells expressing BTK-NanoLuc (Figures S3-S4 and Tables S8-S9). Utilizing established K5-BDP tracers, as described by Promega, we observed strong and reproducible BRET signals at 1 μM tracer concentration, with $IC_{50}$ values ranging from 0.27 ± 0.06 nM to 0.53 ± 0.10 nM and effective displacement of known BTK inhibitor ibrutinib.[4,5] Collectively, these data underscore the tracer system's versatility and scalability across different kinase targets and cellular backgrounds, reinforcing its value for comprehensive target engagement profiling in diverse biological systems.

***Optimization and Validation of Assays for High-Throughput Screening: Key Metrics and Performance Considerations***

The development, optimization, validation, and analysis of assays to assess performance metrics are critical prerequisites for the efficient evaluation of new chemical entities in diverse screening methodologies. These foundational steps have been extensively documented in the literature.[6–8] Prior to full-scale screening, assays must undergo rigorous validation, beginning with the optimization of the dynamic range of signal response and variability. One of the most widely adopted assay quality metrics is the Z' factor, which accounts for both the amplitude and variability of positive and negative responses. This metric is universally used to determine the suitability of an assay for high-throughput screening (HTS). To assess the quality of the assay for high-throughput screening in the absence of test compounds, we computed the Z' factor using the equation below[7,9,10]:

$$Z' = 1 - \frac{3SD_{plateau} + 3SD_{baseline}}{\left|\mu_{plateau} - \mu_{baseline}\right|}$$

Where $SD_{plateau}$ and $SD_{baseline}$, refer to the variability of the positive control signal, which corresponds to the plateau of the assay, and the negative control signal, which represents the background signal. The background signal typically reflects the lowest measurable response, often observed in the absence of specific binding. The terms $\mu_{plateau}$ and $\mu_{baseline}$ represent the mean of the positive and negative control signals, respectively. In TR-FRET and NanoBRET assays, the positive controls correspond to wells containing known binding interactions, while the negative controls contain either no binding activity or a non-specific interaction. A Z' factor greater than 0.5 indicates a robust assay with high signal clarity and low variability between the positive and negative controls, suggesting that the assay is suitable for reliable quantification of binding interactions. Z' values between 0.5 and 1.0 indicate that the assay is effective, with good separation of the signal, while values below 0.5 suggest insufficient assay performance and necessitate optimization. The Z' factor was calculated for each experimental condition, and the filter settings or assay conditions that yielded the highest Z' factor were selected for subsequent analysis and data interpretation.

To ensure reliability in concentration-response assays, key pharmacological parameters—such as $IC_{50}$, $EC_{50}$, and $K_i$—must be characterized using well-established reference compounds. Assay performance measurements (APMs) serve as quantitative metrics for assessing the effectiveness and reliability of biological assays in drug development by measuring signal differences between samples and background controls.

In the absence of test compounds, assay quality for HTS is typically evaluated using the Z' factor. Additional statistical measures, including the signal window, signal-to-blank ratio, signal-to-noise ratio, dynamic range, and limit of detection, also influence assay quality and performance. The signal window, defined as the difference between maximum and minimum signal intensities, is particularly crucial for evaluating assay sensitivity and specificity. Similarly, the signal-to-blank and signal-to-noise ratios, along with the dynamic range and limit of detection, provide insight into assay robustness and reliability. Each of these parameters should be carefully optimized to align with the specific requirements of the assay.

#### *Chemical Stability Analysis of Fluorescent Tracers by LC-MS*

For chemical stability analysis, tracers were incubated at 5 mM in 20 µL complete cell culture medium at 37°C, 5% $CO_2$ for various time points (0, 2, 4, 16, 24 hours), ensuring full protection from light. At each time point, samples were quenched with methanol containing an internal standard (1 mM imipramine) to precipitate proteins, followed by centrifugation and supernatant transfer to LC-MS vials. Samples were then analyzed by reversed-phase liquid chromatography coupled to electrospray ionization mass spectrometry (LC-ESI-MS). Controls included tracer incubated in methanol to distinguish inherent degradation from medium-induced effects, and medium-only blanks to monitor background interference.

#### *Safety Statement*

No unexpected or unusually high safety hazards were encountered.

#### *References*

(6) *Design of Signal Windows in High Throughput Screening Assays for Drug Discovery*. https://doi.org/10.1177/108705719700200306.

(7) Zhang, J.-H.; Chung, T. D. Y.; Oldenburg, K. R. A Simple Statistical Parameter for Use in Evaluation and Validation of High Throughput Screening Assays. *SLAS Discov.* **1999**, *4* (2), 67–73. https://doi.org/10.1177/108705719900400206.

(8) Taylor, P. B.; Stewart, F. P.; Dunnington, D. J.; Quinn, S. T.; Schulz, C. K.; Vaidya, K. S.; Kurali, E.; Lane, T. R.; Xiong, W. C.; Sherrill, T. P.; Snider, J. S.; Terpstra, N. D.; Hertzberg, R. P. Automated Assay Optimization with Integrated Statistics and Smart Robotics. *SLAS Discov.* **2000**, *5* (4), 213–225. https://doi.org/10.1177/108705710000500404.

(9) Bar, H.; Zweifach, A. Z' Does Not Need to Be > 0.5. *Slas Discov.* **2020**, *25* (9), 1000–1008. https://doi.org/10.1177/2472555220942764.

(10) An, W. F.; Tolliday, N. Cell-Based Assays for High-Throughput Screening. *Mol. Biotechnol.* **2010**, *45* (2), 180–186. https://doi.org/10.1007/s12033-010-9251-z.
